# Genome-wide identification, characterization and expression analysis of *CsC3H* gene family in cucumber (*Cucumis sativus* L) under various abiotic stresses

**DOI:** 10.1101/2024.12.23.630068

**Authors:** Saleem Uddin, Sadia Gull, Umer Mahmood

## Abstract

The CCCH zinc finger (C3H) gene family plays a significant role in plant growth, development, and stress response mechanisms. In cucumber (*Cucumis sativus* L), the CsC3H gene family has been implicated in mediating responses to abiotic stresses, though its functional characterization remains underexplored. This study provides a comprehensive genome-wide identification, characterization, and expression analysis of the CsC3H gene family in cucumber, with a focus on their roles in stress tolerance. 38 CsC3H genes have been discovered, and detailed conserved motif and domain analyses revealed key structural features essential for their function. Phylogenetic analysis classified the CsC3H proteins into four distinct subfamilies (CsC3H I–IV), highlighting functional diversification. The gene duplication and expansion analysis indicated that the C3H gene family grew due to both tandem and segmental duplications, with segmental duplications playing a predominant role. qRT-PCR expression profiling revealed widespread expression of CsC3H genes across various cucumber tissues, with distinct differential expression patterns under waterlogging and hormonal treatments (NAA, ETH, MeJA). Notably, CsC3H9 was localized to the nucleus, indicating its potential involvement in regulating cellular processes under stress. These results offer novel information for future studies focused on leveraging genetic innovations to improve stress resilience in cucurbits and other crops while also offering new perspectives on the functional role of CsC3H genes in cucumber.

## 1.1. Introduction

Zinc finger proteins (ZFPs) are a diverse group of sequence-specific transcription factors that contain zinc finger motifs, enabling them to interact with DNA, RNA, and other proteins [1, 2]. Based on the presence of conserved cysteine (C) and histidine (H) residues, ZFPs are categorized into nine subfamilies: C2HC, C2H2, C3H, C4HC3, C3HC4, C4, C2HC5, C6, and C8 [3]. Within these, the C3H-type zinc finger proteins (C3H) are prevalent in eukaryotes and are crucial for the modulating gene expression through establishing direct interactions with mRNA [4, 5]. C3H proteins are characterized by a motif comprising of three cysteine (C) and a single histidine (H) residue, coordinating a zinc ion to build a stable structure which facilitates interactions with nucleic acids and other cellular molecules[6, 7]. The motif sequence is typically characterized as C-X4-15-C-X5-C-X3-H, wherein “X” signifies variable amino acids [8, 9]. A tandem C3H motif (C-X7-8-C-X5-C-X3-H) further divides the C3H family into tandem (TZF) and non-tandem (Non-TZF) zinc finger proteins [10, 11], with TZF proteins being more extensively studied in plants [12].

C3H proteins are involved in several facets of plant development and the response to environmental stresses. For instance, in Arabidopsis, AtC3H59 interacts with the PPPDE protein family to regulate cell cycle progression, apoptosis, and embryogenesis [13]. Other research has demonstrated that non-tandem C3H proteins such as AtC3H17 regulate flowering and senescence [14], while AtC3H2 and AtC3H23 are linked to gibberellin and abscisic acid (ABA) signaling pathways involved in growth and stress responses [15]. C3H proteins from the TZF group also regulate responses to abiotic stress, including salt and drought tolerance. For example, overexpression of AtTZF1 enhances drought tolerance by decreasing transpiration rates [16], while AtTZF1 also regulates cold tolerance by influencing cold-related gene expression [17]. Additionally, C3H genes like AtC3H66, AtC3H20, and AtC3H49 contribute to disease resistance by modulating the plant immune system and enhancing tolerance to abiotic stresses [16, 18].

Cucumber (*Cucumis sativus* L.) is a key crop with significant economic and nutritional value [19]. Gaining insight into the molecular pathways that regulate stress resistance in cucumber is crucial for breeding more resilient cultivars. The C3H gene family has been associated with a range of stress-responsive processes in various plant species, such as cucumber [20]. For example, the C3H-type zinc finger protein CsSEF1 performs a critical function in the formation of somatic embryogenesis [21], and its expression increases in cucumber tissues under defoliation and darkness stress, suggesting a role in stress-related signal transduction [22, 23]. Furthermore, CsSEF1 is upregulated under salinity stress, highlighting its involvement in stress tolerance mechanisms [24–26]. Despite these findings, comprehensive studies on the function of C3H genes in cucumber’s stress resilience and growth and development remain limited.

Cucumber is particularly susceptible to waterlogging stress, which negatively impacts root function and overall plant growth [27]. Waterlogging leads to hypoxia in the root zone, severely affecting plant productivity [28]. Gene expression analyses have shown that waterlogging stress in cucumber induces downregulation of genes involved in carbon metabolism, photosynthesis, and hormone signaling [29]. Several gene families, including ethylene responsive factors (ERFs) and MYB transcription factors, have been implicated in waterlogging responses in cucumber [30, 31], but the specific involvement of C3H zinc finger genes remains underexplored. Some studies suggest that certain C3H genes are upregulated during waterlogging stress, indicating their potential roles in stress adaptation [12]. However, the precise mechanisms through which C3H proteins mediate waterlogging tolerance in cucumber remain unclear.

Genome-wide analysis provides an effective approach for identifying and characterizing gene families across entire genomes [32]. In cucumber, this approach enables the comprehensive cataloging of C3H genes, as well as the analysis of their structural features, evolutionary relationships, and expression patterns under stress conditions [33, 34]. Through the comparative analysis of C3H genes across different species, researchers can uncover the functional diversification and evolutionary patterns of this gene family.

The present study aims to perform genome-wide studies of the C3H zinc finger family in cucumber, with a focus on their roles under waterlogging stress. We identified and systematically analyzed 38 C3H genes in cucumber through various approaches, including phylogenetic analysis, chromosomal mapping, conserved motif identification, gene structure characterization, and assessment of promoter cis-regulatory elements. Additionally, by leveraging RNA-seq data, we investigated the expression patterns of these genes in response to waterlogging stress. Our results offer novel perspective on the biological functional roles of C3H genes in cucumber, laying a foundation for further investigation into their contribution to stress tolerance. These insights could inform future biotechnological approaches to improve waterlogging tolerance in cucumber through targeted breeding and genetic modification.

## 2. Materials and Methods

### 2.3. Phylogenetic Analysis and Identification of *C. sativus* C3H Genes

The C3H domain-containing proteins (PF00642) were identified utilizing the Pfam protein family database (http://pfam.sanger.ac.uk/) and the HMMER tool (http://hmmer.janelia.org/) [35]. We retrieved the protein annotation file from the Ensembl plants database (http://plants.ensembl.org/index). Following, InterPro (http://www.ebi.ac.uk/interpro/) and SMART (http://smart.embl-heidelberg.de/) [36, 37] software were deployed to confirm the reliability of the C3H domain prediction. To examine the biochemical characteristics of CsC3H proteins, the Expasy web platform (https://web.expasy.org/protparam/) was employed. The subcellular localization of the CsC3H genes was inferred through the use of the CELLO2GO online tool (http://cello.life.nctu.edu.tw/) [38]. The complete amino acid sequences of C3H proteins in cucumber were subjected to analysis through multiple sequence alignments conducted using DNAMAN software. For the sequence comparison, ClustalW was utilized (http://www.genome.jp/tools/clustalw/). The maximum likelihood phylogenetic tree was generated using Mega (Version 7.0) software [39]. The designation of each CsC3H gene was determined according to its unique chromosomal location.

### 2.4. Chromosomal locations, gene duplication, and estimation of Ka/Ks values for CsC3Hs

All CsC3Hs were assigned to their corresponding cucumber chromosomes according to their precise locations. The duplication of CsC3H genes in the cucumber genome was detected and illustrated through the application of TBtools software [40]. To delve deeper into the evolutionary divergence of the CsC3H gene duplicates, the nonsynonymous substitution rate (Ka) and synonymous substitution rate (Ks) for each duplicate pair of CsC3Hs were calculated using the Ka/Ks_ Calculator 2.0. This analysis was conducted to assess the divergence between the duplicated CsC3H genes [41]. T= ka/ks 6.1 × 10^-9^ site/year was used to represent the predicted duplication and divergence times MYA (million years ago) for each CsC3H gene utilizing a synonymous mutation rate of λ substitutions per synonymous site per year [42, 43]. The Cucumber genome database provided the CsC3H genes chromosomal position [44, 45].

Moreover, the nomenclature for the *CsC3H* genes was determined by their chromosomal order and their physical locations were plotted using MapDraw.

### 2.5. Collinearity Analysis of CsC3Hs

The chromosomal positions indicated by Ensembl plants were used to map the CsC3Hs to the chromosomes. Gene duplication events were investigated utilizing the Multiple Collinearity Scan toolbox (MCScanX) with default settings [46]. Furthermore, the visualization was generated using Circos (http://circos.ca/) version 0.69, as previously described [47].

### 2.6. Gene structure, protein domains, and motif analysis

The characteristics regarding the isoelectric point (pI), amino acid sequence length (AA), and molecular mass (MW) of each identified amino oxidase protein was obtained from the ProtParam tool available at https://web.expasy.org/protparam [48]. The cucumber GFF3 file was utilized to produce a GFF3 annotation file, which delineated the genomic positions of CsC3Hs along with their structural characteristics. Eventually, the exon-intron organization was visualized using the Gene Structure Display Server (GSDS) (http://gsds.cbi.pku.edu.cn) [49]. The conserved domains of the CsC3H protein sequences were investigated using Pfam (http://pfam.xfam.org/) [50] and the online MEME suite application (http://meme-suite.org) was used to identify the conserved motifs of the cucumber CsC3Hs gene family [51]websites and the resulting files were visualized in TBtools software [52].

### 2.7. *Cis*-regulatory Elements Analysis

To explore the cis-regulatory elements within the promoter region of the CsC3H gene, we retrieved a 2.0 kb sequence upstream of the C3H gene’s initiation codon from the cucumber genome database. This sequence was subsequently uploaded to the PlantCARE platform (http://bioinformatics.psb.ugent.be/webtools/plantcare/html/) for the purpose of predicting potential cis-regulatory elements [53]. The abundance of cis-regulatory elements in the promoter region was then subsequently plotted using TBtools v1.120.

### 2.8. Plant growth and exogenous hormonal treatment

The seeds of *Cucumis sativus* cultivar Chinese Long (CCMC) were incubated on filter paper that had been moistened with distilled water for a duration of 24 hours. Following this, the plants were moved into pots comprising a growth medium composed of peat, vermiculite, and perlite (3:1:1, v/v). When the seedlings developed three to four leaves, they were selected for the stress treatment experiments. These plants were cultivated in a controlled glasshouse with a photoperiod of 14 hours of light and 10 hours of darkness. Daytime temperatures were maintained at 28°C, while nighttime temperatures were kept at 18°C, with relative humidity ranging from 75% to 85%.

For the waterlogging treatment, uniform seedlings, approximately 20–25 days old, were chosen and subjected to waterlogging up to the first cotyledon. Following this, the seedlings were transferred into plastic containers filled with water, ensuring that the water level remained stable throughout the course of the experiment [54]. For the control treatment (0 h), seedlings were placed in identical pots without additional water, following the protocol described by Xu et al. (2023) [55]. Based on preliminary testing with various hormone concentrations, four treatment conditions were established: waterlogging (WL), waterlogging combined with NAA (WNAA), waterlogging combined with ethylene (WETH), and waterlogging combined with methyl jasmonate (WMeJA). To assess the effects of these treatments, Hypocotyls of cucumbers were obtained at 0, 6, 24, 48, and 96 hours after treatment. For each time point and treatment group, three independent biological replicates were obtained, with each replicate consisting of nine hypocotyls. For total RNA extraction and qRT-PCR analysis, the samples were promptly frozen in liquid nitrogen and preserved at -80°C. For exogenous hormone treatments, plantlets at the four-leaf stage were treated with 100 μM/L Naphthalene-acetic acid (NAA), 100 μM/L Ethylene (ETH), and 100 μM/L Methyl jasmonate (MeJA), all applied to the foliage of uniform seedlings.

### 2.9. Transient assay of *CsC3H*

A transient transformation method was employed to investigate the subcellular distribution of the target protein within the epidermal cells of tobacco [56]. The final GFP: CsC3H9 construct was developed by ligating the GFP sequence with the complete open reading frame (ORF) of CsC3H9, which spans 1050 base pairs (primers listed in Table S1). The CsC3H target gene plasmid and the marker plasmid (NAA60-RFP-pCambia2300, used in this experiment) were introduced into Agrobacterium strain GV3101. Following activation and culture expansion of GV3101, the Agrobacterium carrying both the target gene plasmid and the marker plasmid were mixed in equal proportions and re-suspended in tobacco injection buffer to achieve an optical density (OD) value of 1. The mixture was allowed to incubate at room temperature for a period ranging from 1 to 3 hours. Following the incubation, a 1 mL hypodermic syringe was applied to the lower epidermal layer of the tobacco leaves. After 64–88 hours, the epidermis was peeled off, and the samples were stained with DAPI solution. Confocal imaging was conducted using a Leica SP8 confocal microscope (Leica, Germany). The fluorescence of eGFP was captured with an excitation spectrum of 488 nm and an emission range of 493 to 562 nm. In contrast, DAPI staining was visualized using an excitation spectrum of 364 nm and an emission range of 440 to 502 nm.

### 2.10. Profiling of C3H gene expression in *Cucumis sativus*

RNA was isolated from various tissue samples using the Biozol Total RNA Extraction Reagent (Bioer, China). Complementary DNA (cDNA) synthesis was performed with the Reverse Transcriptase cDNA Kit (Jieyi, China) following the provided protocol. The UBIeq gene of cucumber was used as the reference gene for normalization. The expression of C3H genes in Cucumis sativus was assessed using qRT-PCR, utilizing the SYBR Premix Ex Taq II (Takara, Dalian, China) kit and the SYBR Green master mix system (Qiagen) on an ABI PRISM 7500 (Applied Biosystems) platform. Sixteen genes were selected for qRT-PCR analysis based on their expression patterns observed in the heatmap, and the corresponding primer sequences are listed in Table S1. In the experiments, we utilized three biological and technical replicates, following the procedures detailed in our previous research. We quantified relative gene expression using the 2^-ΔΔCT^ method for data analysis [57].

### 2.11. Protein-Protein Interaction and GO ontology analysis

The protein-protein interaction (PPI) networks of rice C3H proteins were uncovered utilizing the STRING database (version 11.0) available at http://string-db.org [58] using default settings. STRING integrates data from various sources, including text mining, experimentally validated interactions, co-expression data, curated repositories, co-occurrence information, and fusion-based approaches. The protein interaction network was visualized using Cytoscape (version 3.7.1) [59], facilitating a comprehensive depiction of the intricate interactions. The functional classification of C3H proteins into biological processes, cellular components, and molecular functions was conducted utilizing the Gene Ontology (GO) database (http://www.geneontology.org/). Fisher’s exact test was utilized to evaluate the enrichment of OsC3H functions in relation to the complete GO database. Functional categories displaying notable enrichment were determined through the application of the Bonferroni correction (Bonferroni52), with a modified significance threshold set at *p* ≤ 0.05 [60].

### 2.12. Statistical Analysis

The statistical analysis was conducted utilizing SPSS software (version 25.0, SPSS Inc., USA), with statistical significance set at *p* ≤ 0.05, corresponding to a 95% confidence level. All data are presented as the mean ± standard deviation (SD) from three independent replicates for each evaluated parameter. Data visualization was conducted with GraphPad Prism 7.0 (GraphPad Software, Inc., La Jolla, CA, USA). Heatmaps of RNA-seq data were generated using TBtools [40], with Log2-transformed RPKM values used for normalization.

## 3. Results

3.1. Identification and Characterization of *CsC3H* transcription factor family members in *Cucumis sativus*

This research identified 38 members of the C3H gene family in the cucumber genome utilizing HMMER 3.0 software, with the analysis anchored on the C3H domain (PF00642). These 38 C3H genes were mapped onto chromosomes and designated as CsC3H1 through CsC3H38 in accordance with their chromosomal positions (Tab 1). The CsC3H genes’ coding sequences varied in length, with CsC3H1 spanning from 5725813 bp and CsC3H9 extending to 19941834 bp. The amino acid profile of CsC3H proteins ranged from 156 in CsC3H3 to 958 in CsC3H11. While four CsC3H proteins contained fewer than 300 amino acids, the majority (34 proteins) exceeded 300 amino acids in length. The isoelectric points (pI) of the CsC3H proteins varied, with CsC3H13 exhibiting the highest pI of 9.35 and CsC3H1 showing the lowest at 4.96. Molecular weights varied from 17.87 kDa for CsC3H3 to 113.02 kDa for CsC3H11. The grand average hydropathy (GRAVY) values for all CsC3H proteins were negative, ranging from -0.157 for CsC3H15 to -1.244 for CsC3H11. Furthermore, most CsC3H proteins are situated within the nucleus, while CsC3H4 is situated in the plasma membrane. Additional details regarding the chromosomal locations, locus IDs, molecular weights and subcellular localizations of the identified C3H zinc finger proteins in *Cucumis sativus* are provided in Table 1.

**Table 1:**
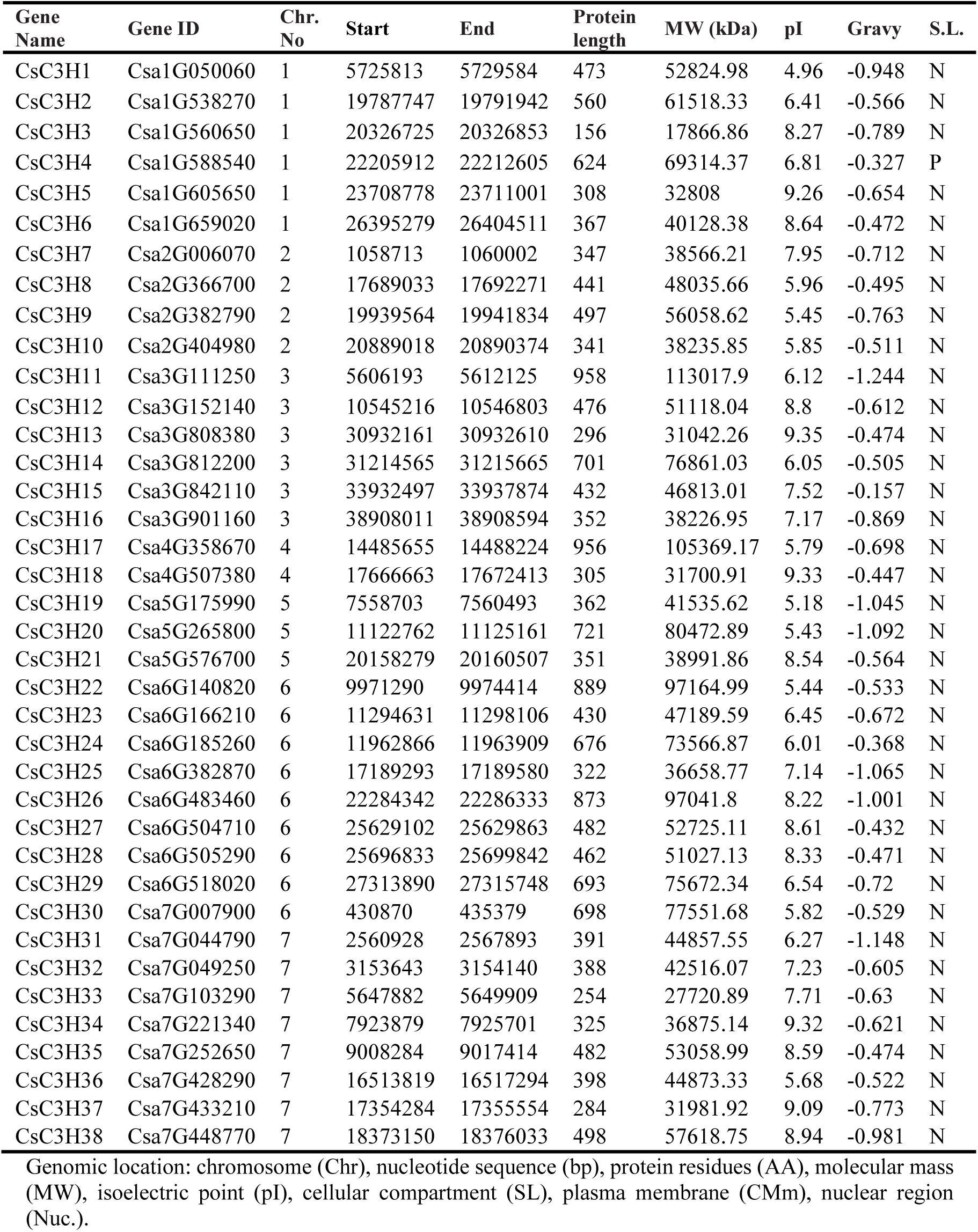
Properties of genes and proteins in the CsC3Hs family of *C. sativus*.

| Gene Name | Gene ID | Chr. No | Start | End | Protein length | MW (kDa) | pI | Gravy | S.L. |
| --- | --- | --- | --- | --- | --- | --- | --- | --- | --- |
| CsC3H1 | Csa1G050060 | 1 | 5725813 | 5729584 | 473 | 52824.98 | 4.96 | -0.948 | N |
| CsC3H2 | Csa1G538270 | 1 | 19787747 | 19791942 | 560 | 61518.33 | 6.41 | -0.566 | N |
| CsC3H3 | Csa1G560650 | 1 | 20326725 | 20326853 | 156 | 17866.86 | 8.27 | -0.789 | N |
| CsC3H4 | Csa1G588540 | 1 | 22205912 | 22212605 | 624 | 69314.37 | 6.81 | -0.327 | P |
| CsC3H5 | Csa1G605650 | 1 | 23708778 | 23711001 | 308 | 32808 | 9.26 | -0.654 | N |
| CsC3H6 | Csa1G659020 | 1 | 26395279 | 26404511 | 367 | 40128.38 | 8.64 | -0.472 | N |
| CsC3H7 | Csa2G006070 | 2 | 1058713 | 1060002 | 347 | 38566.21 | 7.95 | -0.712 | N |
| CsC3H8 | Csa2G366700 | 2 | 17689033 | 17692271 | 441 | 48035.66 | 5.96 | -0.495 | N |
| CsC3H9 | Csa2G382790 | 2 | 19939564 | 19941834 | 497 | 56058.62 | 5.45 | -0.763 | N |
| CsC3H10 | Csa2G404980 | 2 | 20889018 | 20890374 | 341 | 38235.85 | 5.85 | -0.511 | N |
| CsC3H11 | Csa3G111250 | 3 | 5606193 | 5612125 | 958 | 113017.9 | 6.12 | -1.244 | N |
| CsC3H12 | Csa3G152140 | 3 | 10545216 | 10546803 | 476 | 51118.04 | 8.8 | -0.612 | N |
| CsC3H13 | Csa3G808380 | 3 | 30932161 | 30932610 | 296 | 31042.26 | 9.35 | -0.474 | N |
| CsC3H14 | Csa3G812200 | 3 | 31214565 | 31215665 | 701 | 76861.03 | 6.05 | -0.505 | N |
| CsC3H15 | Csa3G842110 | 3 | 33932497 | 33937874 | 432 | 46813.01 | 7.52 | -0.157 | N |
| CsC3H16 | Csa3G901160 | 3 | 38908011 | 38908594 | 352 | 38226.95 | 7.17 | -0.869 | N |
| CsC3H17 | Csa4G358670 | 4 | 14485655 | 14488224 | 956 | 105369.17 | 5.79 | -0.698 | N |
| CsC3H18 | Csa4G507380 | 4 | 17666663 | 17672413 | 305 | 31700.91 | 9.33 | -0.447 | N |
| CsC3H19 | Csa5G175990 | 5 | 7558703 | 7560493 | 362 | 41535.62 | 5.18 | -1.045 | N |
| CsC3H20 | Csa5G265800 | 5 | 11122762 | 11125161 | 721 | 80472.89 | 5.43 | -1.092 | N |
| CsC3H21 | Csa5G576700 | 5 | 20158279 | 20160507 | 351 | 38991.86 | 8.54 | -0.564 | N |
| CsC3H22 | Csa6G140820 | 6 | 9971290 | 9974414 | 889 | 97164.99 | 5.44 | -0.533 | N |
| CsC3H23 | Csa6G166210 | 6 | 11294631 | 11298106 | 430 | 47189.59 | 6.45 | -0.672 | N |
| CsC3H24 | Csa6G185260 | 6 | 11962866 | 11963909 | 676 | 73566.87 | 6.01 | -0.368 | N |
| CsC3H25 | Csa6G382870 | 6 | 17189293 | 17189580 | 322 | 36658.77 | 7.14 | -1.065 | N |
| CsC3H26 | Csa6G483460 | 6 | 22284342 | 22286333 | 873 | 97041.8 | 8.22 | -1.001 | N |
| CsC3H27 | Csa6G504710 | 6 | 25629102 | 25629863 | 482 | 52725.11 | 8.61 | -0.432 | N |
| CsC3H28 | Csa6G505290 | 6 | 25696833 | 25699842 | 462 | 51027.13 | 8.33 | -0.471 | N |
| CsC3H29 | Csa6G518020 | 6 | 27313890 | 27315748 | 693 | 75672.34 | 6.54 | -0.72 | N |
| CsC3H30 | Csa7G007900 | 6 | 430870 | 435379 | 698 | 77551.68 | 5.82 | -0.529 | N |
| CsC3H31 | Csa7G044790 | 7 | 2560928 | 2567893 | 391 | 44857.55 | 6.27 | -1.148 | N |
| CsC3H32 | Csa7G049250 | 7 | 3153643 | 3154140 | 388 | 42516.07 | 7.23 | -0.605 | N |
| CsC3H33 | Csa7G103290 | 7 | 5647882 | 5649909 | 254 | 27720.89 | 7.71 | -0.63 | N |
| CsC3H34 | Csa7G221340 | 7 | 7923879 | 7925701 | 325 | 36875.14 | 9.32 | -0.621 | N |
| CsC3H35 | Csa7G252650 | 7 | 9008284 | 9017414 | 482 | 53058.99 | 8.59 | -0.474 | N |
| CsC3H36 | Csa7G428290 | 7 | 16513819 | 16517294 | 398 | 44873.33 | 5.68 | -0.522 | N |
| CsC3H37 | Csa7G433210 | 7 | 17354284 | 17355554 | 284 | 31981.92 | 9.09 | -0.773 | N |
| CsC3H38 | Csa7G448770 | 7 | 18373150 | 18376033 | 498 | 57618.75 | 8.94 | -0.981 | N |

Genomic location: chromosome (Chr), nucleotide sequence (bp), protein residues (AA), molecular mass (MW), isoelectric point (pI), cellular compartment (SL), plasma membrane (CMm), nuclear region (Nuc.).

3.2. Evolutionary relationships of *CsC3H* gene family in *C. sativus*

A maximum likelihood Phylogenetic tree was generated employing the entire amino acid full-length C3H protein sequences of each member of *Cucumis sativus*, *Oryza sativa* and *Arabidopsis thaliana* to explore insightful knowledge of the evolutionary connections between C3H zinc finger genes. The complete amino acid sequences of 38 CsC3H, 52 AtC3H, and 57 OsC3H genes were used to generate the phylogenetic tree. Furthermore, the CsC3H protein family was categorized into four distinct subgroups: CLADE I, CLADE II, CLADE III, and CLADE IV. The quantity of CsC3H zinc finger proteins within respective subgroup was quantified. Of these, the CLADE II and CLADE III groups emerged as the largest, each comprising 12 genes, followed by CLADE IV with 11 genes, and CLADE I, which contained only 3 genes (Figure 1). The distribution of C3H zinc finger genes across these subgroups was notably uneven.

**Fig. 1.**
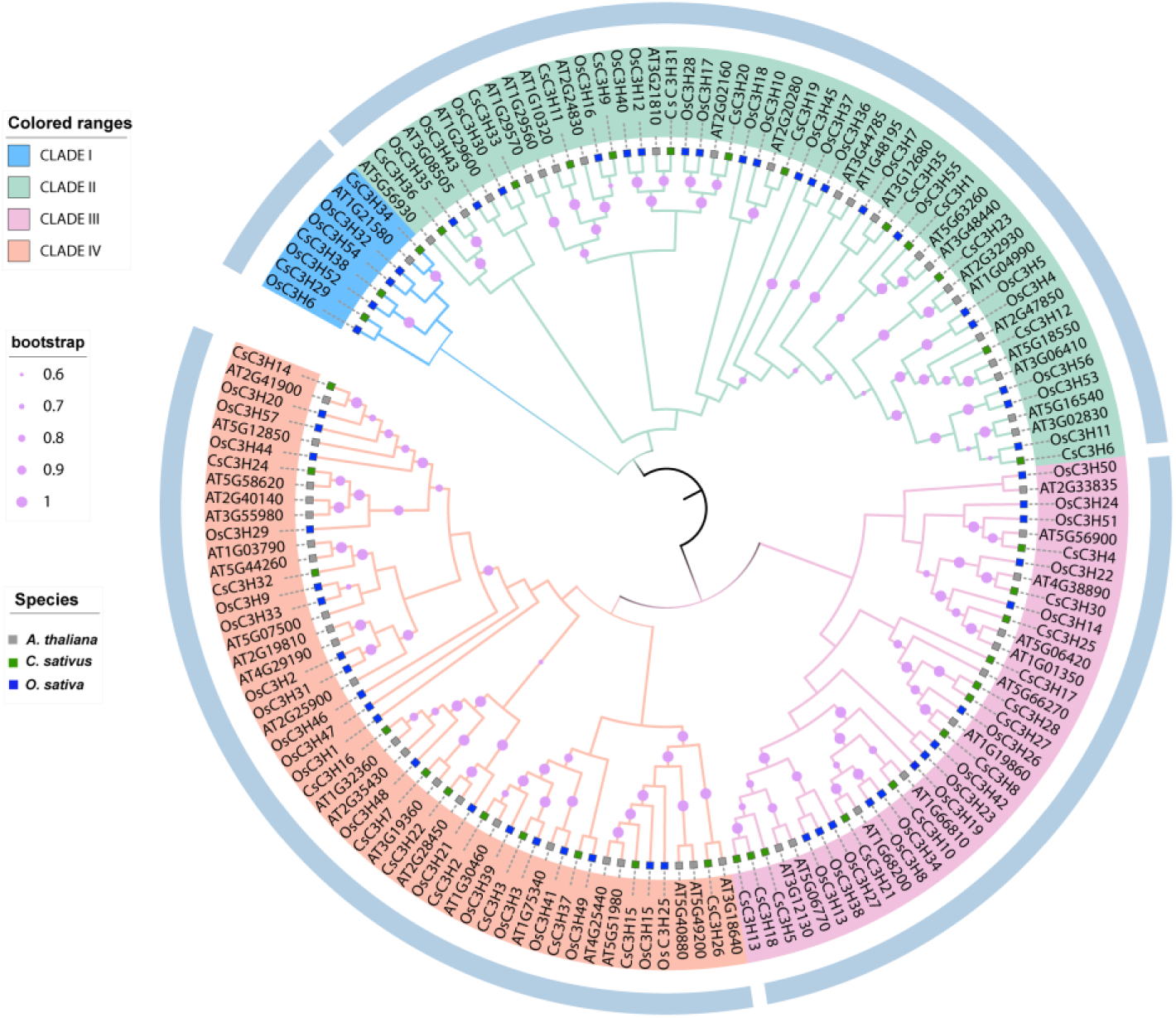
Phylogenetic overview of the *CsC3H* gene family in *C. sativus*. A tree was constructed using maximum likelihood based on the protein sequences of *CsC3H* genes from cucumber and selected plant species (*Arabidopsis thaliana* and *Oryza sativa*). Bootstrap values (1,000 replicates) are shown at the nodes. The tree reveals distinct clustering patterns, providing insights into the phylogenetic relationships and potential efficient diversification of the C3H gene family.

### 3.3. Analysis of genomic localization and evolutionary characteristics of the *CsC3H* gene family in *C. sativus*

Based on the genome annotation of *C. sativus*, the distribution of C3H zinc finger genes across its chromosomes was characterized. A total of 38 C3H zinc finger genes were detected and positioned across chromosomes 1 through 7. These 38 *CsC3H* genes exhibited an uneven distribution, and the number of genes located on each chromosome was not correlated with the chromosome’s size. The largest chromosome, Chr06, harbored the highest number of *CsC3H* genes, with nine genes mapped to it, followed by Chr07, which contained eight genes. Six genes were positioned on both Chr01 and Chr03, while Chr02 contained four genes, Chr05 had three, and the smallest chromosome, Chr04, had two genes (Figure 2). Gene duplication, involving segmental and tandem duplications, is prevalent in plant genomes and is regarded as a significant factor in genome evolution, leading to substantial gene family expansion in plants. Duplicated genes provide a foundation for generating new genetic variation. Consequently, we detected tandem duplication events through multiple collinearity analyses. Four gene pairs *CsC3H1/CsC3H5*, *CsC3H11/CsC3H15*, *CsC3H27/CsC3H28*, *CsC3H26/CsC3H29*, were discovered as tandem duplicated genes and resided on chromosomes Chr01, Chr03 and Chr06. Tandem duplication events were not identified on Chr02, Chr04, Chr05, and Chr07. These results reveal that tandem duplication events have been essential in promoting the diversification of C3H genes in *C. sativus.* Additionally, segmental duplication events of *CsC3H* genes were also identified, suggesting their role in the evolutionary development of this gene family. A total of seventeen pairs of genes resulting from segmental duplications were detected, distributed across all seven chromosomes. Among the 38 *CsC3H* genes, 37% (14 genes) were involved in segmental duplications, leading to the identification of 30 segmentally duplicated gene pairs (Figure 3). Furthermore, we performed a phylogenetic analysis to examine the nonsynonymous substitution rate (Ka), synonymous substitution rate (Ks), and the Ka/Ks ratio, along with investigating the chromosomal localization of the *CsC3H* genes. The findings indicated that the *CsC3H* gene duplication events likely occurred between 40.2 and 4.6 million years ago (MYA). Notably, the majority of the Ka/Ks ratios for the gene pairs were found to be below 1.0, suggesting that these gene pairs underwent purifying selection following duplication (see supplementary table 1).

**Fig. 2.**
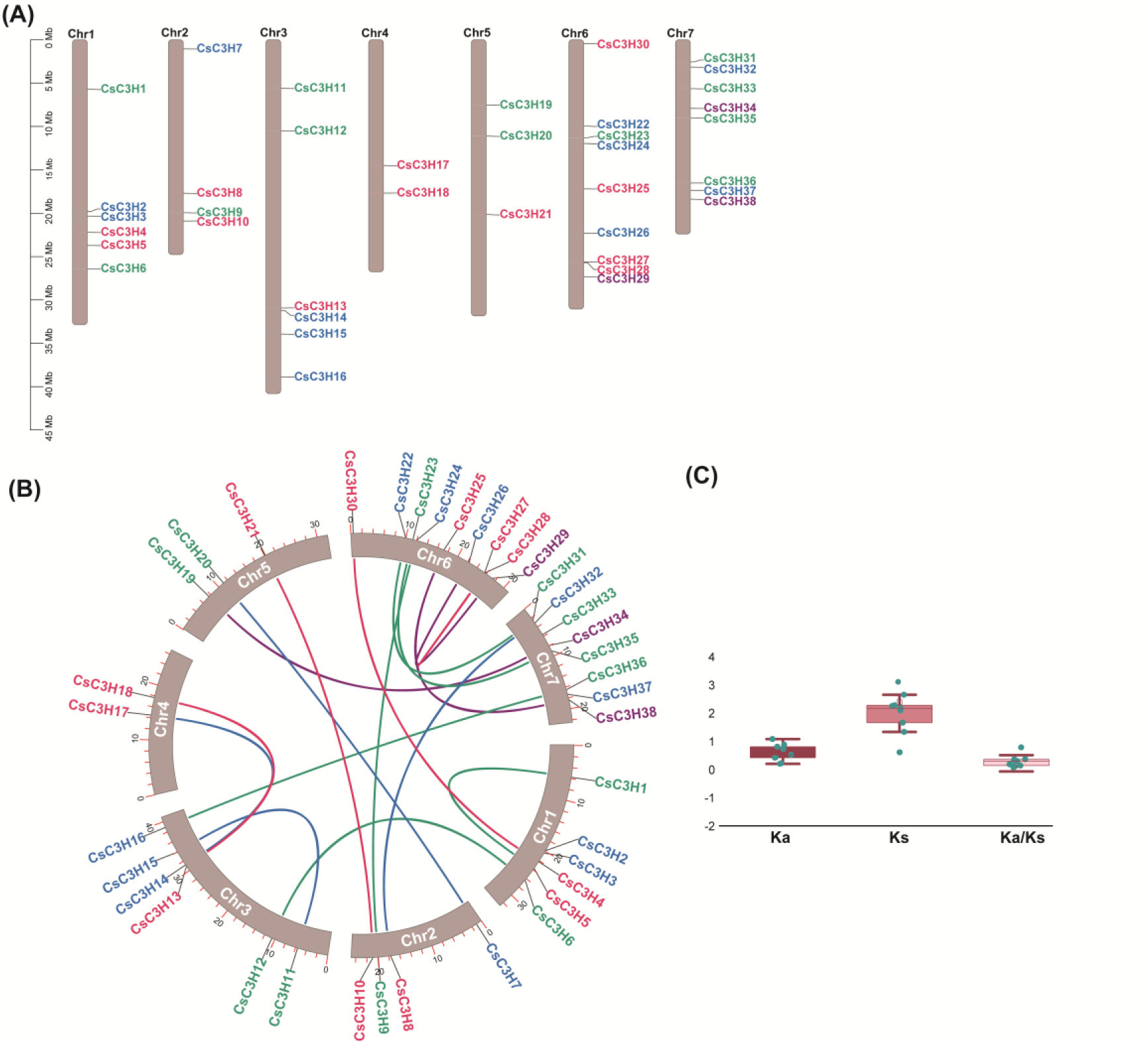
(A) Chromosomal distribution of *CsC3H* genes across *C. sativus* chromosomes (Chr1–Chr7), showing gene locations and densities. **(B)** Collinearity analysis of *CsC3H* genes within *C. sativus*, highlighting duplication events and evolutionary relationships. **(C)** Ka/Ks ratios of *CsC3H* gene pairs, indicating selective pressures acting on gene evolution.

**Fig. 3.**
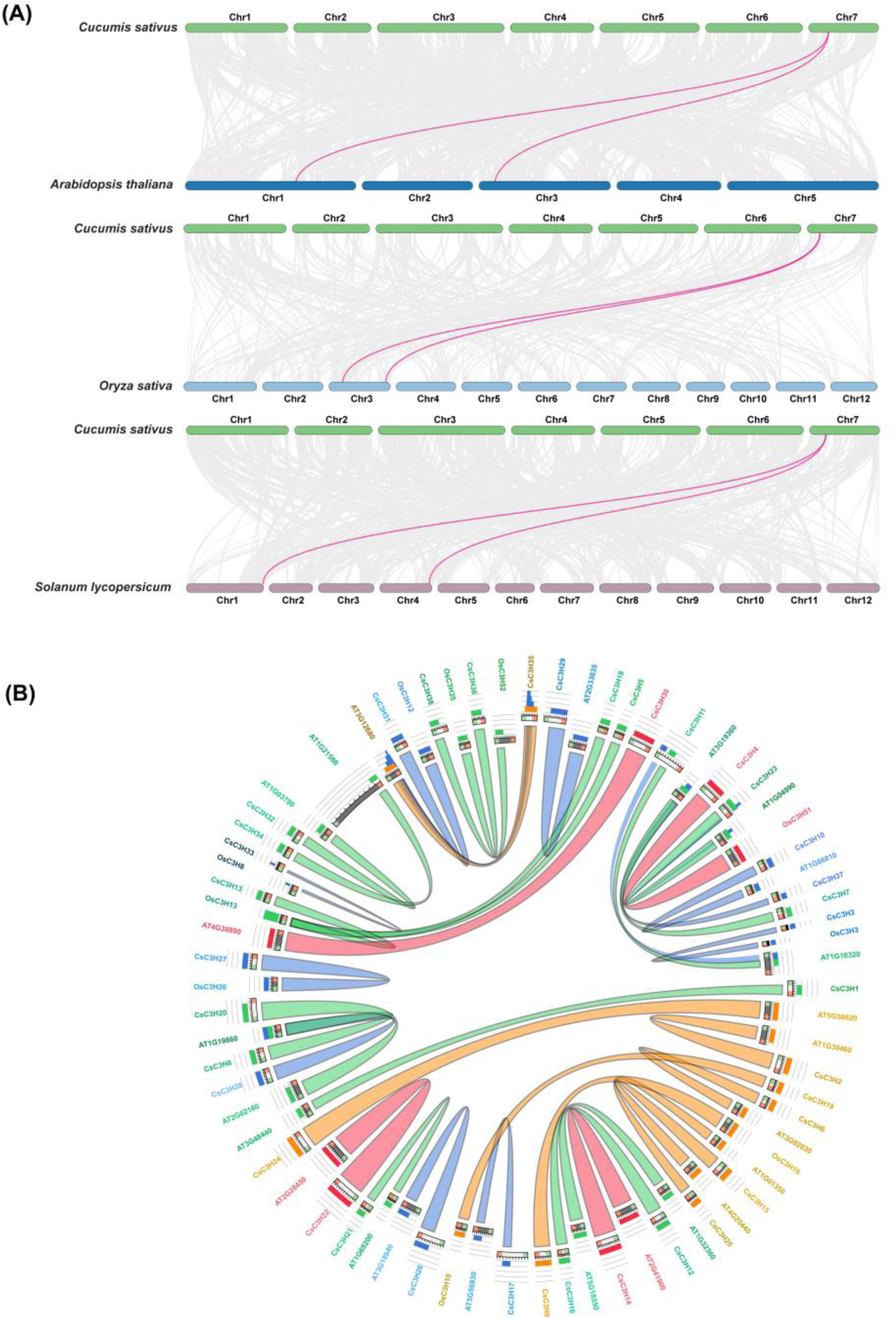
Comparative synteny analysis of the *C3H* gene. **(A)** Synteny analysis of the *C3H* gene family in cucumber. Syntenic regions between cucumber and related species (Arabidopsis, rice) are displayed, with conserved gene pairs highlighted by lines connecting orthologs. The analysis reveals chromosomal rearrangements and gene conservation across species, providing understanding the evolutionary processes of the *CsC3H* gene family **(B)** Circos plot illustrates syntenic relationships, showing high chromosomal conservation and significant similarity across species (bitscore > 0.75).

### 3.4. Comparative Synteny of *CsC3H* genes across Cucumber, Arabidopsis, Rice and tomato genomes

We further investigated collinearity and relationships between *Cucumis sativus CsC3H* genes and corresponding genes from *Solanum lycopersicum* and *Arabidopsis thaliana* to categorize homologous genes. In total 6 homologous gene pairs were exhibited an association between *C. sativus, Arabidopsis thaliana and S. lycopersicum.* Collinearity relationships were identified with 4 genes between *C. sativus* and Arabidopsis genes and 2 *C. sativus* genes between *S. lycopersicum* genes (Figure 3).In addition, we exhibited the syntenic similarities between *C. sativus*, *A. thaliana* and *O. sativa CsC3H* genes through Circos. Our findings reveal that all three genomes exhibited a significant degree of synteny with the genome of *C. sativus*. The resulting synteny blocks displayed a strong chromosomal conservation among the *CsC3H* genes of *C. sativus*, *A. thaliana*, and *O. sativa*. Highest similarities were observed between *CsC3H4/OsC3H*, *CsC3H30/AT4G38890*, *CsC3H22/AT2G28450* and *CsC3H14/AT2G41900*.

### 3.5. Comprehensive assessment of conserved motifs, domains and Gene structure of the *CsC3H* gene family in *C. sativus*

The structures of *CsC3H* genes in *C. sativus* were evaluated to understand the evolution of gene diversification. Results show that the intron/exon number of *CsC3H* genes varies from 2–16, but each subfamily was relatively conserved in gene structure. Genes that are closely associated, especially those demonstrating collinearity, show considerable resemblance in their exon-intron arrangements, with the primary variation observed being in the intron length. Specifically, the collinear gene pairs *CsC3H14/CsC3H24, CsC3H6/CsC3H12, and CsC3H10/CsC3H21* display remarkable similarities in their motifs, protein domains, and exon/intron configurations. Nevertheless, differences in the length of exons and introns result in substantial variations in the overall gene length (Figure 4A). There are significant differences in the motif composition and exon-intron architecture across different groups, suggesting functional divergence within each subgroup of the *C. sativus CsC3H* gene family. A comprehensive analysis of the conserved motifs within this gene family identified ten distinct motifs (Figure 4B). Motifs 1 and 2 were recognized as highly conserved regions, present in nearly all *CsC3H* genes, indicating their critical importance in the functionality of CsC3H proteins. Additional motifs, such as motifs 5, 7, 9, and 10, were found exclusively in Class I, implying that *CsC3H* genes within the same class share similar motif structures, while distinct variations were observed between different classes. This suggests that *CsC3H* genes within a single class may perform analogous functions, and certain motifs may be pivotal for specific functional roles (Figure 4B). All *CsC3H* genes analyzed were found to contain the YTH, MISS, PAT1, DEC-1_N, zf-C2H2_jaz, WD40, Nup160, G-patch, and TIM domains (Figure 4C).

**Fig. 4.**
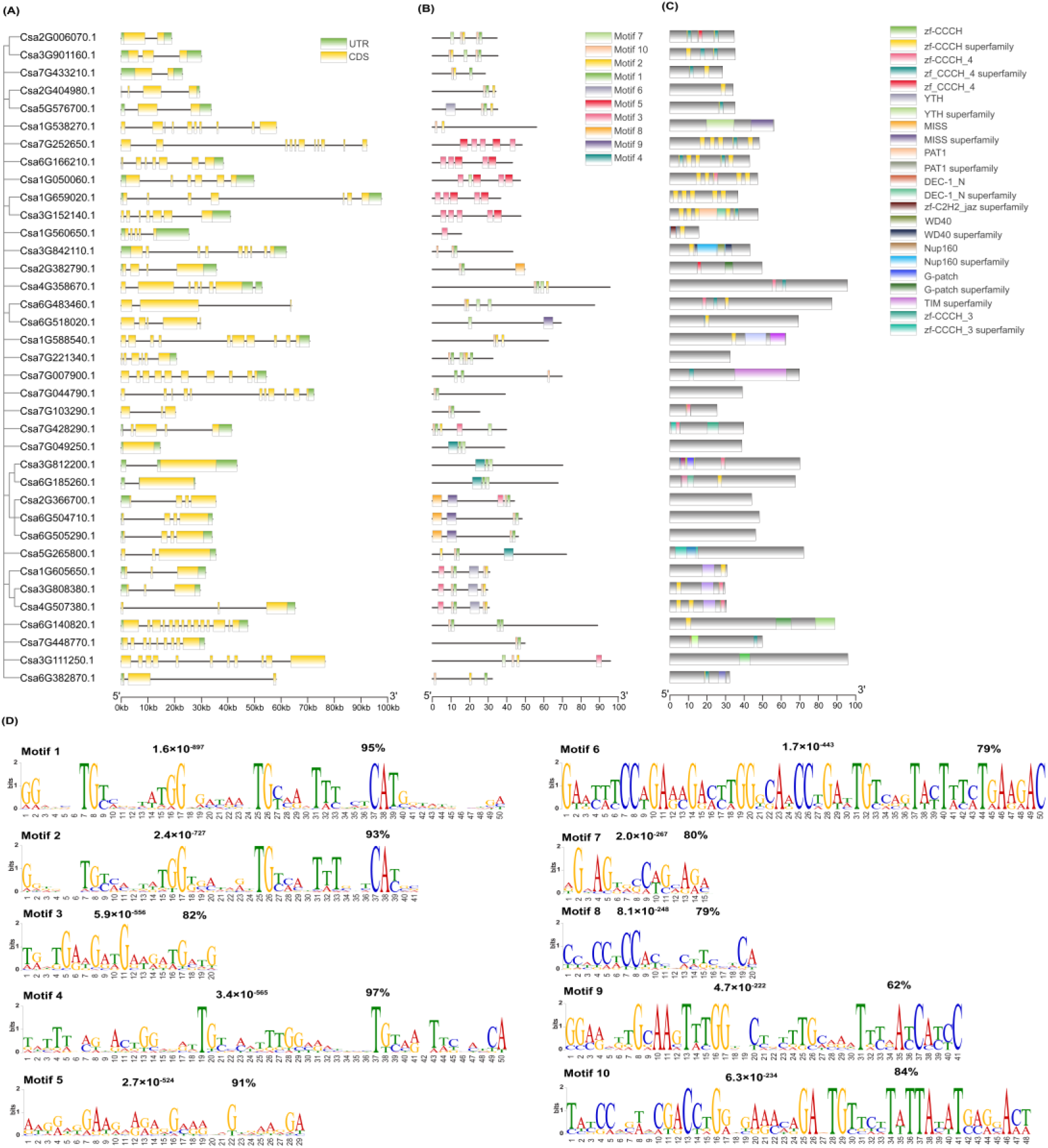
Analysis of conserved motifs, domains and gene structure of *CsC3H* gene family members in *C. sativus*. **(A)** Schematic representation of conserved motifs identified in CsC3H proteins, with each motif displayed in a distinct color. **(B)** Conserved domain structures, highlighting the *CsC3H* domain superfamily and other domains within *CsC3H* members. **(C)** Exon-intron structures of *CsC3H* genes, with coding sequences (CDS) and untranslated regions (UTRs) differentiated by color. Scale bars indicate sequence and structural alignment. **(D)** Sequence logos of conserved motifs within CsC3H proteins, illustrating nucleotide frequency at each position within the motif. Bit scores indicate the conservation level of individual bases across the aligned sequences.

3.6. Analysis of cis-regulatory elements within the *CsC3H* gene family in *C. sativus*

A comprehensive examination of the cis-regulatory motifs and expression profiles of the *CsC3H* gene family in *C. sativus* revealed a complex regulatory landscape, offering insights into their functional dynamics and potential regulatory mechanisms. After analyzing the 2000 bp upstream promoter regions, a total of 720 cis-regulatory elements were identified. These elements were classified into four distinct categories: 221 light-responsive elements, 297 hormone-responsive elements, 78 elements associated with plant growth and development, and 124 elements linked to environmental stress responses (Figure 5). *Cis*-regulatory elements associated with stress, such as MYB-binding sites and dehydration-responsive elements, were particularly abundant in *CsC3H2* and *CsC3H17* (7), whereas several other genes, including *CsC3H6*, *CsC3H18*, and *CsC3H31*, harbored only one. Hormone-responsive cis-regulatory elements, including ABRE and TCA-elements, were most abundant in *CsC3H22* (26), while *CsC3H7* and *CsC3H33* exhibited the lowest counts. Plant growth and development-related elements peaked in *CsC3H17* (8), with minimal representation in multiple genes. *cis*-acting regulatory elements associated with Light-responsive regulation, such as G-box and GT1-motifs, were most frequent in *CsC3H26* (18), while *CsC3H3*, *CsC3H19*, and *CsC3H25* contained only one. The comprehensive analysis of cis-regulatory elements within the promoter regions of the *CsC3H* gene family offers valuable insights into their potential regulatory functions and expression profiles, thereby deepening our comprehension of their functional mechanisms.

**Fig. 5.**
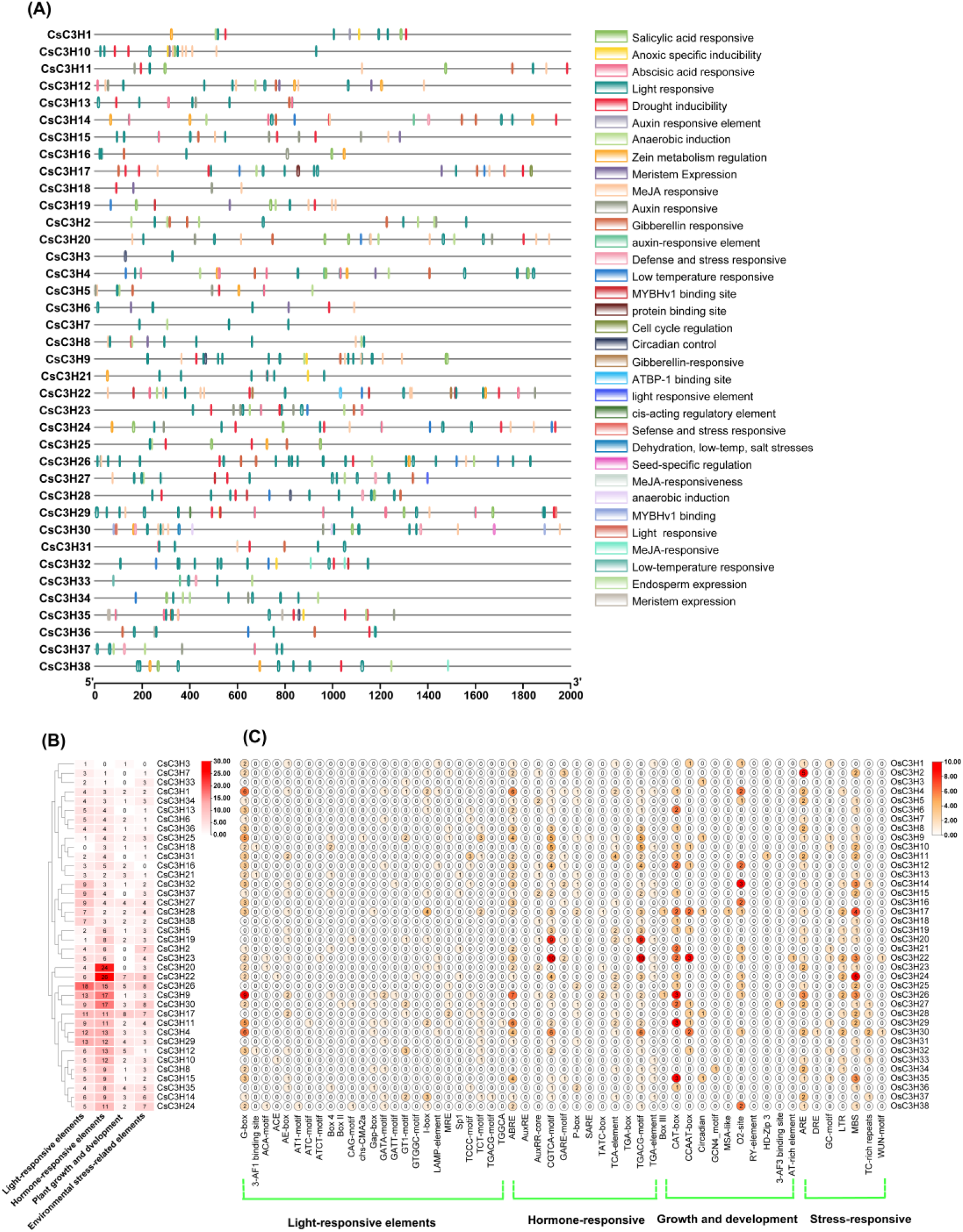
*Cis*-acting regulatory element study of *CsC3H* genes in cucumber. **(A)** Distribution of *cis*-regulatory elements in the promoter regions of *CsC3H* genes, highlighting elements related to stress response, phytohormone signaling, and developmental processes. **(B)** Relative frequency of key cis-elements associated with abiotic stress (e.g., ABRE, DRE) and hormone responses (e.g., ERE, AuxRE) across *CsC3H* gene promoters, illustrating potential regulatory roles in stress adaptation and growth modulation.

### 3.7. Protein-Protein Interaction Network Analysis of C3H genes in cucumber

We constructed the protein interaction network of the cucumber *CsC3H* gene family members utilizing the STRING online tool, referencing the cucumber protein database. This analysis revealed potential interactions among 31 of the family members (Figure 6). In contrast, no interaction relationships were detected for the remaining six proteins of the cucumber C3H family. These findings indicate that the members of the cucumber C3H gene family likely engage in intricate multigenic interactions rather than direct protein-protein interactions, suggesting a broader, network-based regulatory mechanism rather than direct protein-to-protein connections. The protein functions identified in *C. sativus* (Csa) offer significant insights into the molecular pathways governing different cellular processes. For instance, Csa_6G453810 is involved in RNA decapping and RNA processing, playing a crucial role in controlling mRNA stability and gene expression. Similarly, Csa_6G367180 contributes to stress responses, particularly those involving high light intensity and hydrogen peroxide, while also participating in D-amino acid metabolism. Csa_5G412760 is implicated in metabolic processes with hydrolase activity, likely supporting key biochemical reactions in the cytosol. Csa_1G000770, a transcription factor, takes part in controlling gene expression at the transcriptional and post-transcriptional phases, highlighting its essential role in nuclear processes. Proteins such as Csa_6G148350 and Csa_4G642540 are linked to acyl-carrier-protein biosynthesis, playing pivotal roles in lipid metabolism, cell wall modification, and energy storage. Csa_4G165870 and Csa_4G165890 participate in RNA catabolism and processing through their roles in the exosome complex, ensuring proper RNA turnover and stability. Csa_3G601030 contributes to phosphoinositide metabolism, which is crucial for cellular signaling and membrane dynamics, while Csa_6G401320 is involved in pseudouridine synthesis, modifying RNA for enhanced stability and function. Csa_1G478590 is essential for coenzyme A biosynthesis and nucleotide metabolism, processes critical for energy production and cellular metabolic regulation. Csa_5G622860 and Csa_6G405350 are involved in RNA decapping and degradation, key steps in mRNA regulation in the cytoplasm. Csa_5G175730 plays a substantial function in RNA surveillance and the degradation of defective mRNA, ensuring the maintenance of RNA quality. Csa_6G146430 is involved in various plant developmental processes, including floral organ formation and seed germination, and contributes to chromatin modification, influencing gene expression during development. Finally, Csa_3G627150 is involved in protein targeting to mitochondria, which is essential for mitochondrial function and energy production. Csa_5G155570, another transcription factor, regulates gene expression by binding DNA, while Csa_2G374620 contributes to RNA surveillance, splicing, and protein maturation, ensuring proper RNA and protein function within the cell. These proteins form a network that regulates key processes like RNA metabolism, stress response, and plant development, supporting homeostasis and function in *C. sativus*.

**Fig. 6.**
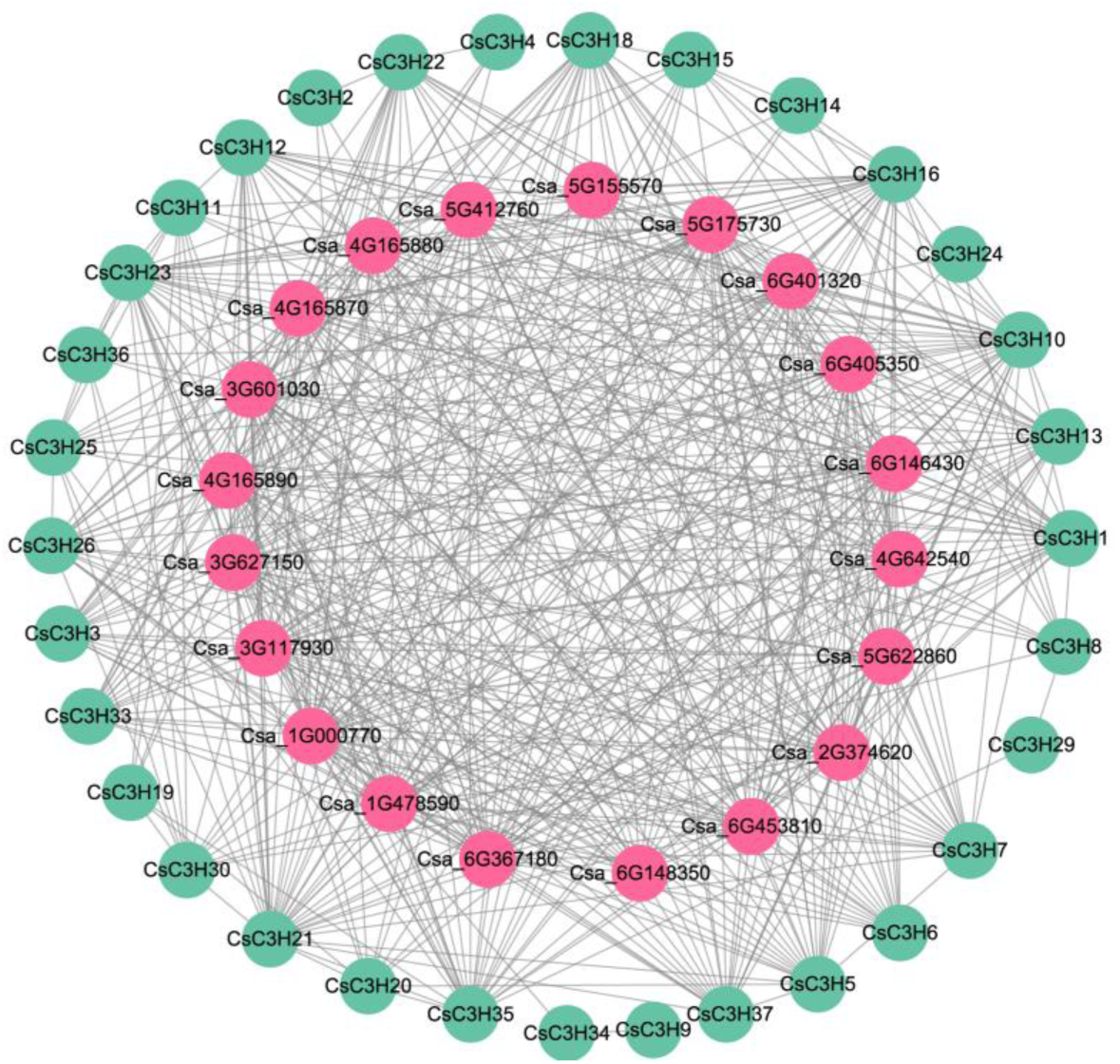
Interaction networks of the *CsC3H* gene family in *C. sativus*. **(A)** The protein-protein interaction (PPI) network illustrates interactions among *C. sativus CsC3H* proteins, where nodes represent CsC3H proteins and edges indicate predicted interactions. **(B)** miRNA target network showing predicted miRNAs and their corresponding *CsC3H* gene targets, with miRNAs indicated by nodes linked to target genes through directed edges.

### 3.9. Gene ontology (GO) analysis of *CsC3H* genes

A thorough Gene Ontology (GO) analysis was performed to explore and elucidate the functional implications of *CsC3H* genes. The results indicated that the *CsC3H* gene family is implicated in various regulatory functions across molecular activities, cellular structures, and biological phenomena (Fig. 7). These genes are associated with several biological processes, including the modulation of transcription, DNA-dependent mechanisms, nucleic acid phosphodiester bond cleavage, metabolic pathways, and transport functions. Additionally, *CsC3H* genes are involved in cellular components, particularly within the plastid and nucleus. They also contribute to protein interactions and exhibit transcription factor activity with specificity for DNA sequence recognition (Fig. 7).

**Fig. 7.**
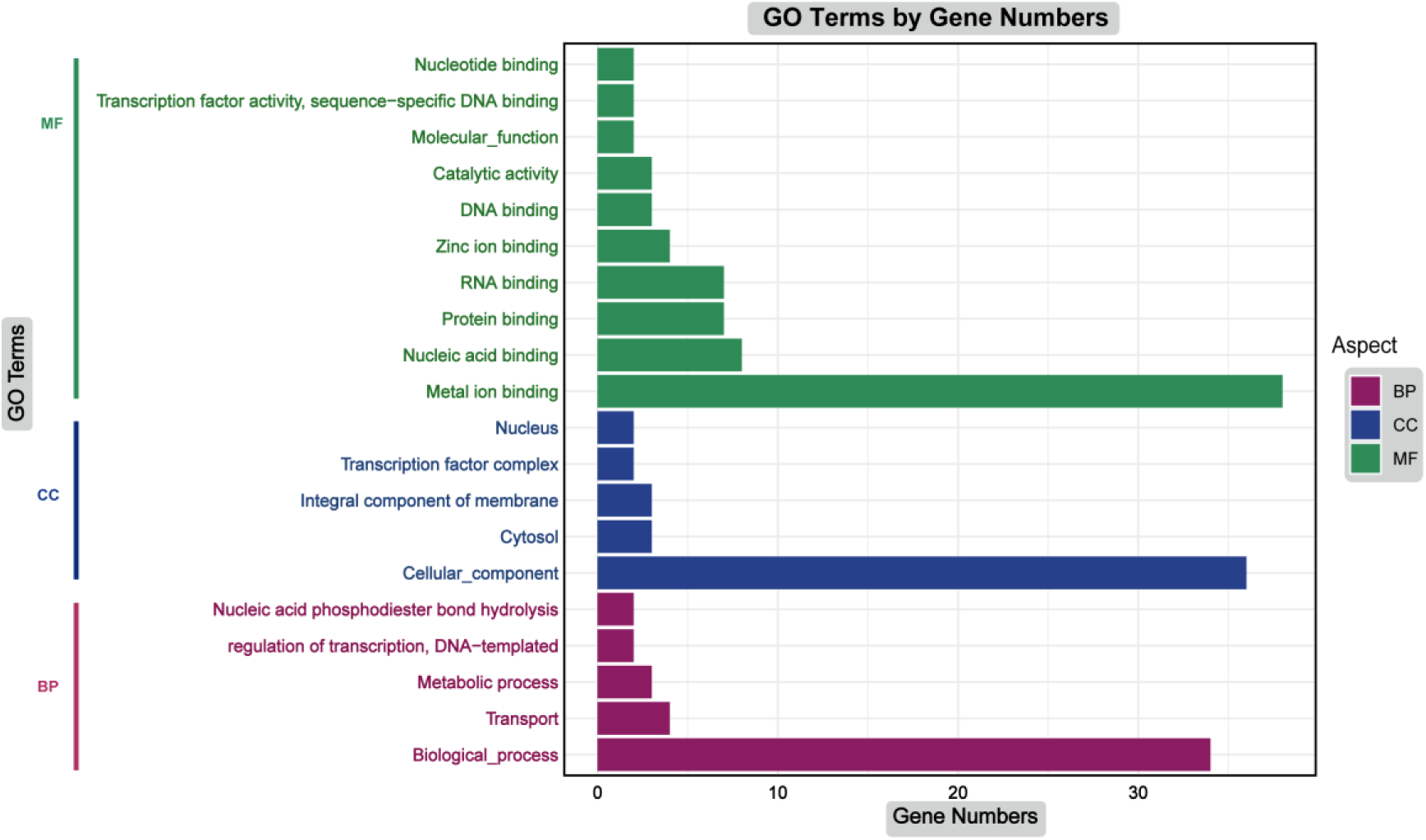
Gene Ontology (GO) study of the *CsC3H* genes in *C. sativus*. The data is presented in three distinct categories: molecular function (MF), cellular component (CC), and biological process (BP).

### 3.10. Effect of waterlogging stress and exogenous hormones on the Phenotype of *C. sativus*

To examine the effects of waterlogging stress independently and in combination with three exogenous hormone treatments, phenotypic assessments were conducted on cucumber hypocotyls over a 96-hour period. From 0 hours to 3 days, the control (0 days), waterlogging (WL), waterlogging + NAA, waterlogging + ETH, and waterlogging + WMeJA treatments exhibited relatively similar phenotypes at 3 days. In contrast to plants subjected solely to waterlogging (WL) stress for 48 hours, the formation of adventitious roots (ARs) on the hypocotyl was significantly lower compared to plants treated with WL combined with either NAA or ETH. No ARs were observed on plants treated with waterlogging combined with MeJA. After 4 days of WL stress, treatment with WL and NAA resulted in a substantial increase in AR formation, followed by WL and ETH treatments, both of which induced more ARs than WL treatment alone. Consistently, no ARs were detected on MeJA-treated plants (Figure 8A). The number of adventitious roots (ARs) was significantly higher in WNNA- and WETH-treated plants compared to those treated with WL, while no ARs were observed in WMeJA-treated plants under 96 hours of treatment (Figure 8B).

**Fig. 8.**
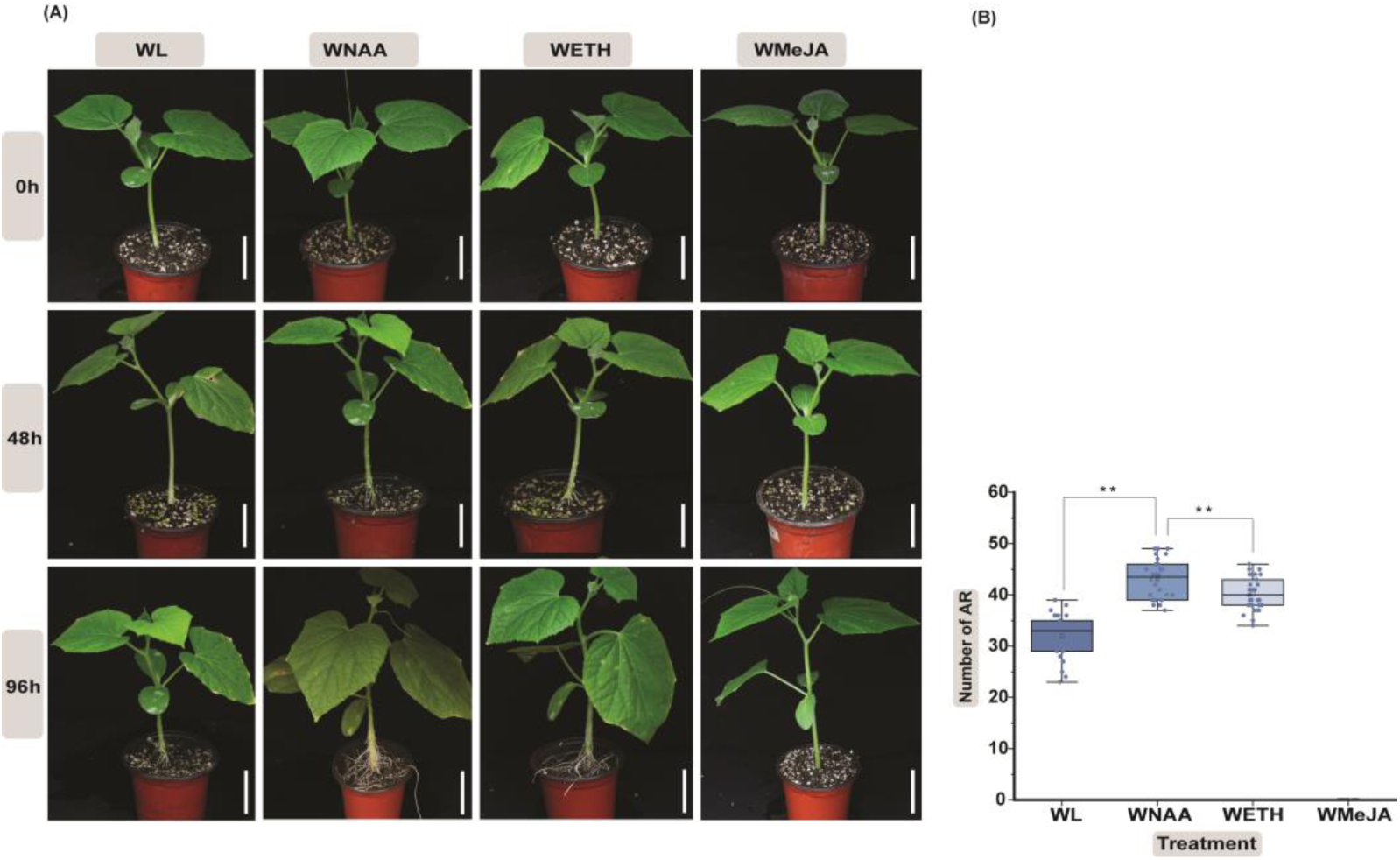
Phenotypic adaptations and statistics in response to waterlogging stress and combination with Waterlogging+NAA, Waterlogging+ETH and Waterlogging+WMeJA hormonal treatments. **(A)** The response of waterlogging and waterlogging+NAA, waterlogging+ETH and waterlogging+WMeJA hormones treatment. **(B)** Capacity of adventitious roots (AR) formation under various treatments, ** *p* < 0.01, *t-*test, (*n*=3).

### 3.11. Expression patterns of some *CsC3H* genes under waterlogging stress combined with hormones treatment

Our study measured the expression profile of 12 representative *CsC3H* genes that were significantly up-regulated in response to waterlogging stress combined with NAA, ETH, and MeJA treatment by qRT-PCR for various stress periods, 0, 6, 12, 24, 48 and 96 hours (Figure 9). As predicted, the *CsC3H* genes exhibited diverse expression profiles during treatment with waterlogging and various hormones. Under waterlogging treatment, *CsC3H9* and *CsC3H13* genes showed a positive response with a significantly increasing expression profile, while *CsC3H2, CsC3H4, CsC3H7, CsC3H11, CsC3H12,* and *CsC3H15* exhibit significantly lower expression under detected abiotic stresses. Whereas, 5 *CsC3H* genes (*CsC3H2, 3, 12, 13,* and *14*) showed higher sensitivity to NAA treatment, on the other hand, 4 *CsC3H* genes (*CsC3H4, 7, 11,* and *15*) showed decrease expression level to NAA treatment. Moreover, *CsC3H6, CsC3H12* and *CsC3H15* showed high sensitivity under ETH treatment. The majority of *CsC3H* genes demonstrated substantial down-regulation upon MeJA exposure, with the notable exceptions of *CsC3H5*, *CsC3H7*, and *CsC3H11*, indicating their potential involvement in the plant’s response to waterlogging conditions. Twelve *CsC3H* genes responded to ETH in varying patterns. In general, most of the *CsC3H* genes were sensitive to various hormone treatments. Furthermore, 3 *CsC3H* genes (*CsC3H*6, 12 and 13) were significantly up-regulated more than two times under NAA and ETH treatments and down-regulated more than two times in MeJA and waterlogging treatment. However, *CsC3H6* was up-regulated in all four treatments, it is proposed that it could function as a pleiotropic regulator, modulating responses to various biotic stresses and hormonal treatments.

**Fig. 9.**
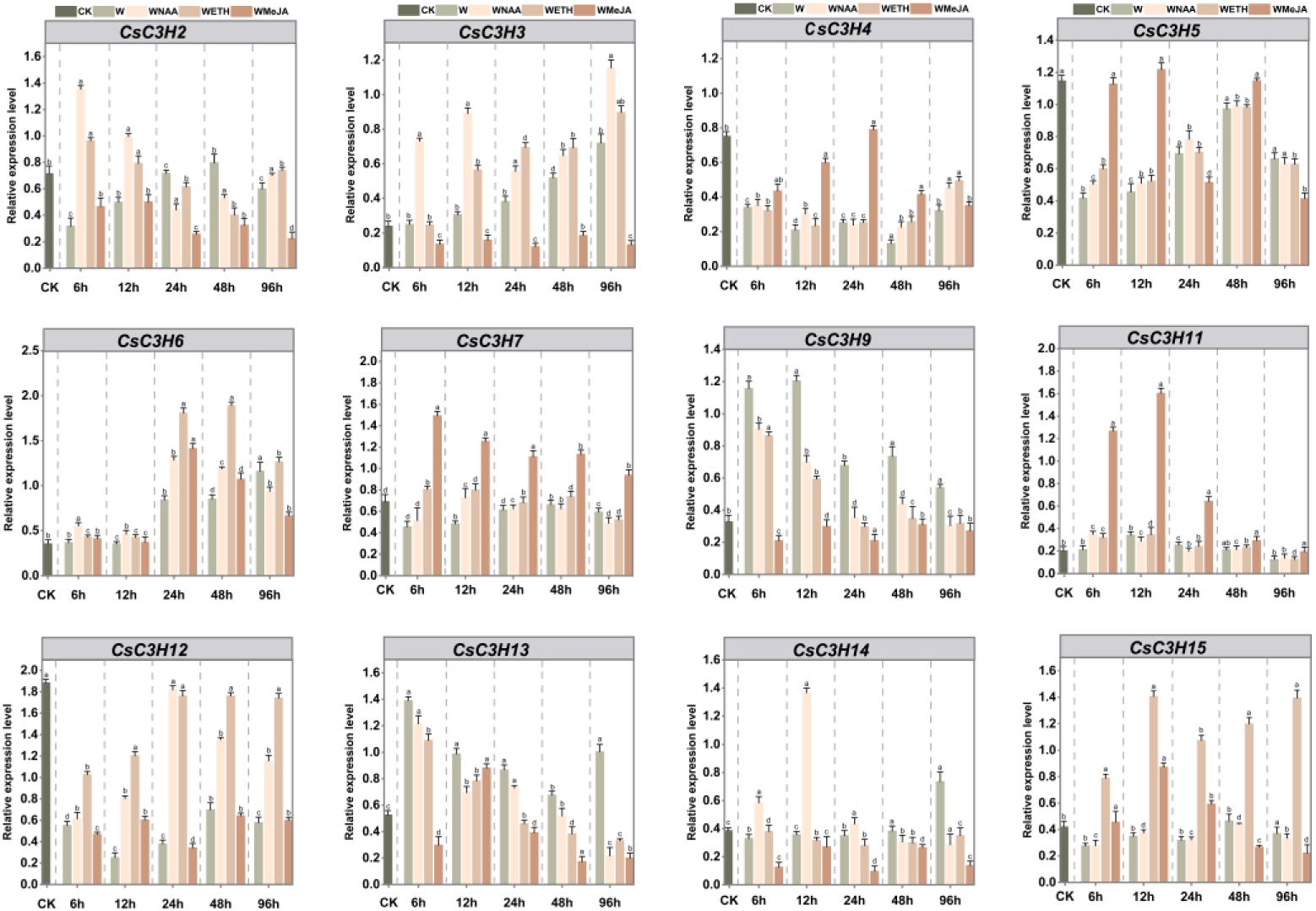
Relative expression profile of *CsC3H* genes at different time points in response to waterlogging combined with hormonal treatments. Different time points are shown on the x-axis, while fold changes are displayed on the y-axis. QRT-PCR determined the relative expression profiles of 12 genes. The distinct letters (a, b, c, etc.) on the bars represent statistically significant changes. The results reflect three independent trials (n=3, mean ±SD, *\*p* < 0.05).

### 3.12. Transcriptional expression pattern analysis of cucumber *CsC3H* genes under waterlogging stress and different tissues

To explore the broader role of *CsC3Hs*, transcriptomic data were employed to analyze its expression patterns. The waterlogging stress, in combination with three exogenous hormones, NAA, ETH and MeJA, was investigated in time courses 0, 6, 12, 24, 48, and 96 hours. The heatmap results projected the expression level of *CsC3Hs* revealed significant differences across the stress period and exogenous hormone treatments compared with control. Among the 38 *CsC3Hs,* 10 genes showed limited expression throughout the whole stress period, though the expression profile of 8 genes experienced minor variations. For instance, *CsC3H17* and *CsC3H32* did not undergo notable changes during the control and waterlogging stress period, and their expression profile was down-regulated during the exogenous hormone treatments. Throughout the entirety of hormonal treatments and abiotic stress time course, the expression levels of maximum *CsC3Hs* were increased, including *CsC3H2/3/4/5/6/7/9/11/12/14/15* and *CsC3H*24 demonstrated a consistent upward trend during different time course. For example, *CsC3H3, CsC3H6* and *CsC3H15* expression levels were constantly up-regulated during the hormonal treatments in contrast to the control plants (Figure 10A).

**Fig. 10.**
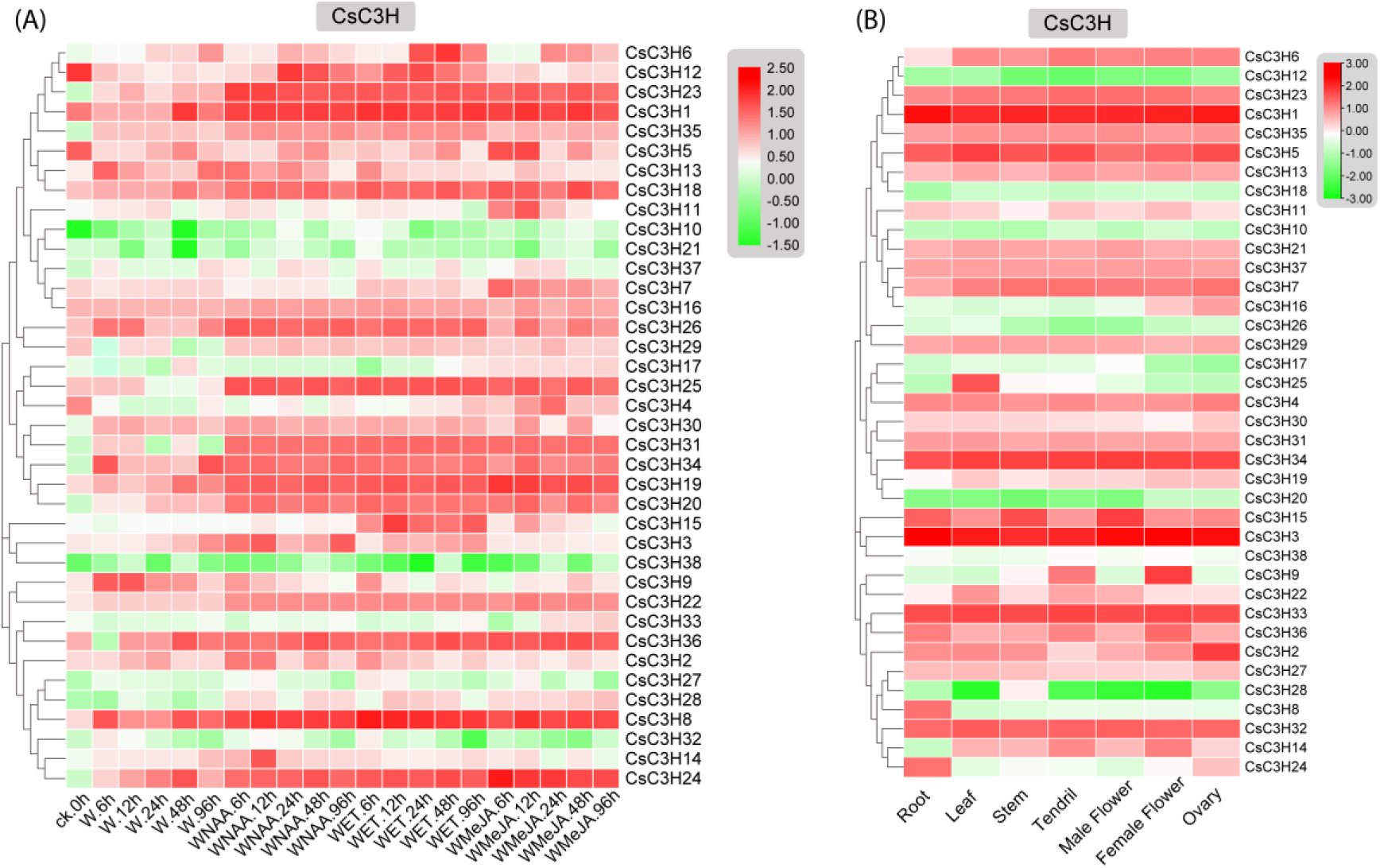
Heatmaps of *CsC3H* genes in post exogenous hormones treatment combine with waterlogging stress and different tissues of *C. sativus* **(A)** The expression profile of *CsC3H* genes under various exogenous hormones treatments combine with waterlogging stress. **(B)** The expression pattern of *CsC3H* genes across various tissues. The log^2^ transformation method was used to normalize and convert the RFKM values, as illustrated on the color scale bars.

The transcriptional activity of *CsC3H* genes was assessed across various plant tissues, including the root, leaf, stem, tendrils, male flower, female flower, and ovary. The heatmap analysis revealed that 38 *CsC3H* genes demonstrated tissue-specific expression profiles in cucumber. Predominantly, these genes were expressed in tendrils, leaves, roots, stems, and ovaries of female flowers, and male flowers, indicating their significant role in diverse physiological and developmental processes (Figure 10B). Specifically, *CsC3H2*, *CsC3H7* and *CsC3H5* are dominantly expressed in ovaries, suggesting its involvement in seed development. Whereas, *CsC3H22, CsC3H25,* and *CsC3H32 are* positively expressed in leaf tissues. In addition, *CsC3H*9 and *CsC3H*36 exhibit high expression in female flower tissues, signifying their positive role in the development of flowers and fruit. Furthermore, seven genes (*CsC3H10*, *CsC3H12*, *CsC3H17*, *CsC3H18*, *CsC3H20*, *CsC3H28* and *CsC3H38*) down-regulated in all tissues. In developmental processes such as flowering and fruit, down-regulation of some genes can be vital for transitioning from the vegetative to the reproductive phase. Particularly in our study, *CsC3H12, CsC3H17* and *CsC3H28* significantly down-regulated in female flower developmental phase, which can hypothesize that these genes are involved in some process of fruit development.

To evaluate the tissue-specific expression and in depth relative expression patterns assessment of *CsC3Hs* genes, we carried out qRT-PCR assays on a subset of selected genes (Figure 11). The results revealed that two genes, *CsC3H9* and *CsC3H14* exhibited higher expression in female flower followed by tendril tissues in contrast with other tissues which exhibit low expression. The higher expression in female flowers specifies the vital role of *CsC3Hs* genes in the formation of flower and fruit. The expression profile of *CsC3H5* was similar to that of *CsC3H22* and *CsC3H25.* Its expression level in the leaf tissues was significantly greater in contrast with the tendrils, root and stem tissues. High expressions of these genes in the leaf tissues indicate their potential profound role in leaf development in *C. sativus*. The expression trend of *CsC3H8* and *CsC3H24* was higher in the roots exceeds than in the leaf and ovary. Moreover, *CsC3H2* was significantly expressed in ovary, followed by female flower and leaf tissues, whereas *CsC3H15* predominantly expressed in the male flowers followed by stem and root tissues.

**Fig. 11.**
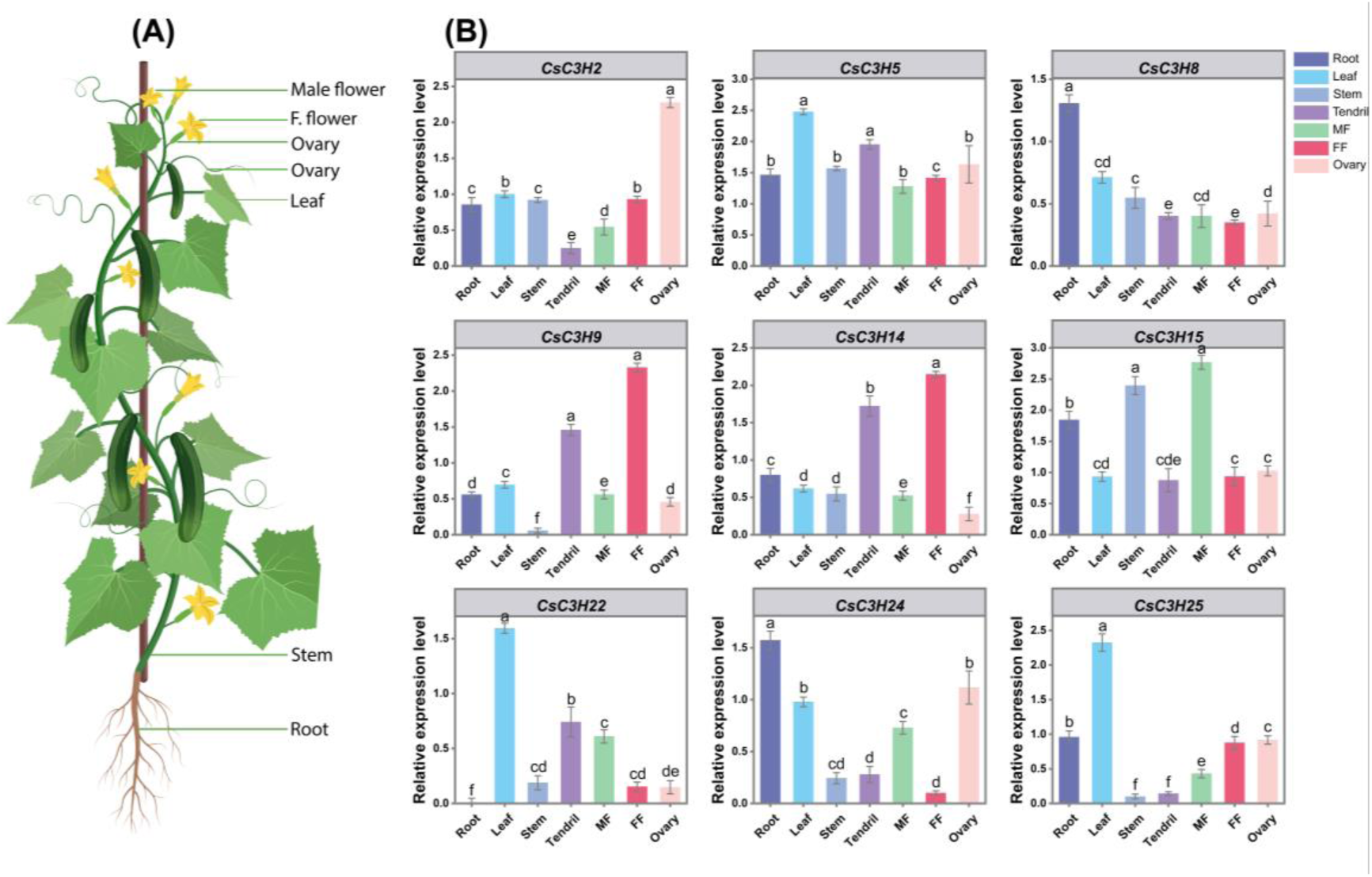
Tissue expression pattern analysis of *C. sativus CsC3H* genes. Different colors depicts various tissue types. “MF” denotes the male flower, while “FF” represents the female flower. The error bars indicate the mean ± standard deviation from three independent replicates. Distinct lowercase letters above the bars signify a statistically significant difference (*p* < 0.05).

### 3.13. Analysis of cucumber CsC3H protein Subcellular localization

The majority of CsC3H protein is located in the nucleus, as mentioned in Table 1; however, to better comprehend the role of the *CsC3H* gene family, in silico prediction of cucumber CsC3H protein through subcellular localization of the resulting GFP-tagged fusion protein was investigated, the *35S::CsC3H9-GFP* and *35S::D53*-*RFP* plasmids were transiently expressed into *N. benthamiana* epidermal cells for fluorescence microscopy. For nuclear localization, *35S::D53-RFP* plasmid assisted as a positive control, which expressed a red fluorescent protein precisely targeted to the nucleus. After introducing the vector into tobacco leaves containing the nuclear localization marker, fluorescence signals were detected. The result of subcellular localization was consistent with the predicted results and showed that *CsC3H9* is specifically expressed in the nucleus (Figure 12).

**Fig. 12.**
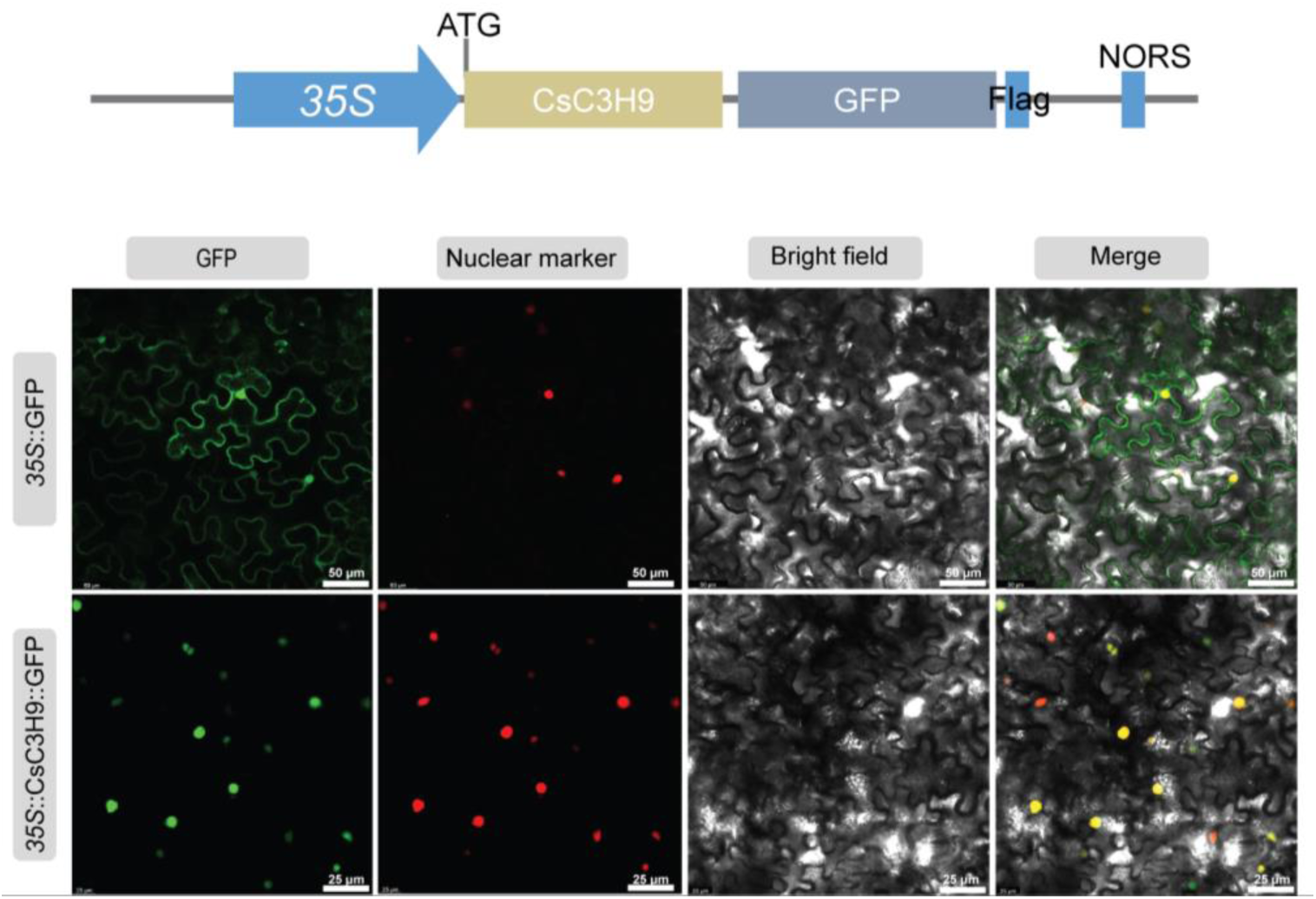
Subcellular localization of Cucumber *CsC3H* was analyzed—the GFP-fused in *N. benthamiana* epidermal cells for fluorescence microscopy. The figure from left to right depicts GFP channel, nuclear marker, bright field, and merged image. 35S::GFP is utilized as a control, 35::D53-RFP is a nuclear marker. Scale bar = 50 and 25 µm.

## 4. Discussion

Cucumbers (*Cucumis sativus* L.) are an important horticultural crop, widely cultivated for their nutritional value. Its genome serves as a valuable resource for understanding plant quality improvement and stress resistance mechanisms [44]. Zinc finger proteins, a key family of transcription factors (TFs), play a crucial role in regulating plant growth, gene regulation, and responses to environmental stress, with the C3H-type zinc finger family drawing particular attention for its role in cell differentiation, stress response, and protein interactions [61–63]. Although much is known about other zinc finger types, such as C2H2 and C3H, the C3H family remains under-researched, especially in species like cucumber. A genome-wide examination of gene families across the entire genome provides crucial insights into the regulatory mechanisms governing plant biological processes, establishing a foundation for identifying potential candidates for deeper functional studies.

With the advancement of in silico tools, it is now possible to conduct genome-wide predictions of gene families. Using these tools, the cucumber genome was analyzed to identify 38 C3H zinc finger genes. These 38 genes were identified with high confidence, contributing to the understanding of C3H gene dynamics in cucumber. Interestingly, the number of C3H proteins in cucumber was found to be lower than that in species comprising *Arabidopsis thaliana* (67), *Oryza sativa* (68), *Zea mays* (68), *Vitis vinifera* (69), *Populus* (91), *Panicum virgatum* (103), and *Brassica rapa* (103) [4, 5, 20, 34, 64–67]. This suggests that factors like species origin and genome size do not directly correlate with the number of C3H genes. The 38 CsC3H proteins exhibit diverse isoelectric points (pI), a characteristic linked to solubility, subcellular localization, and protein interactions. In general, proteins found in the cytoplasm exhibit low isoelectric points (pI), typically below 4.96, whereas those localized in the nucleus are characterized by more neutral pI values, ranging from 4.96 to 9.35 [68, 69]. The variation in isoelectric points, along with differences in gene length and motif patterns, indicates that the C3H genes in cucumber are contributed in a broad spectrum of biological functions, potentially controlling number of developmental attributes in the plant.

Phylogenetic investigations are essential for understanding the evolutionary relationships between species [70]. Phylogenetic analysis revealed that the C3H genes from cucumber, *Oryza sativa*, and *Arabidopsis thaliana* could be classified into four distinct groups (Figure 1). This classification aligns with groupings found in other plant species, such as *Cicer arietinum*, *Phaseolus vulgaris*, *Glycine max*, and *Hordeum vulgare* [71–73]. Each group contained C3H genes from all species studied, with cucumber C3H genes (*CsC3H*) sharing significant homology with their counterparts in other species, particularly *Oryza sativa* and *Arabidopsis thaliana* [74]. These results suggest that a common ancestral gene likely gave rise to pairs of orthologs, reflecting a conserved evolutionary lineage of C3H genes across these plants.

Chromosomal distribution and duplication patterns of *CsC3H* genes in cucumber provide insight into their evolutionary dynamics. The distribution of the 38 *CsC3H* genes on the chromosome is not uniform, with the highest concentration on chromosomes 6 and 7, which may be influenced by local genomic architecture or selective pressures, as seen in other plants gene families (Figure 2) [75]. Tandem duplications on chromosomes 1, 3, and 6 contributed significantly to the evolution of the gene family, while the absence of duplications on chromosomes 2, 4, 5, and 7 could reflect selective constraints. Segmental duplications, accounting for 17 events across all chromosomes, were also key to CsC3H gene expansion, with 37% of the *CsC3H* genes involved in these events. The Ka/Ks analysis indicated that most gene pairs are under purifying selection, suggesting their evolutionary conservation [76]. The duplication events, which occurred between 40.2 and 4.6 million years ago, imply substantial diversification of the *CsC3H* gene family. Tandem and segmental duplications have extensively contributed in the enlargement and diversification of the *CsC3H* gene family, thereby laying a solid groundwork for future investigations into their functional contributions, especially concerning stress resistance and adaptability.

A more detailed analysis of the gene structure and motif constitution within the cucumber C3H gene family demonstrated that *CsC3Hs* within the same cluster exhibited similar intron/exon configurations, intron phases, and conserved functional domains. This observation indicates a preservation of function across evolutionary processes. In comparison with other species, such as *Populus trichocarpa* (15 conserved domains), *Arabidopsis* (13), *Medicago truncatula* (8), *Vitis vinifera* (6), *Zea mays* (10) and *Citrus mandarin* (16), the cucumber C3H family contains 9 conserved domains, which suggests functional diversification within the family [4, 20, 66, 67, 77, 78]. In various biological processes such as the transport of molecules, signal transduction, recognition of RNA and DNA, RNA binding, and interactions between proteins, several conserved domains such as ANK, RRM, WD-40, KH, and others play crucial roles [79]. Additionally, cucumber C3H members possess several unique conserved domains, including the CCCH, YTH, MISS, PAT1, DEC-1_N, zf-C2H2_jaz, WD40, Nup160, G-patch, and TIM superfamilies, suggesting their involvement in a wide array of biological processes (Figure 4) [12, 80–82]. A number of *cis*-acting regulatory elements identified in the promoters of CsC3H genes (Figure 5) offer valuable insights into the complex roles of C3H proteins in plant development and their participation in defense mechanisms against both biotic and abiotic stresses [83–85]. In this study, we identified four distinct types of *cis*-regulatory elements: those responsive to light, hormones, growth and regulatory processes, as well as stress-related elements. These elements were found to be closely linked with particular *CsC3H* genes.

CCCH zinc-finger proteins are involved in various functions related to plant adaptation to waterlogged stress. Hypoxic conditions in rice induce the transcription of OsCCCH-Zn-1, OsCCCH-Zn-2, and OsCCCH-Zn-3. The expression of OsCCCH-Zn-1 is significantly upregulated in response to flooding stress and ABA treatment, while remaining unchanged under H2O2, cold, or salinity stress conditions. This suggests that OsCCCH-Zn-1 potentially specifically control flooding stress responses via the ABA pathway, with minimal involvement in other abiotic stress responses [65]. Additional investigation is required to fully elucidate the mechanisms by which CCCH zinc-finger proteins facilitate plant adaptation to flooding.

The CCCH zinc-finger proteins in plants play a crucial role in modulating stress responses that are mediated by hormones [86]. These proteins are often induced by hormones such as jasmonic acid (JA), ABA, and gibberellin (GA), they play critical roles in a variety of hormone-driven signaling pathways [87, 88]. Environmental stresses, including drought, waterlogging, salinity, heat, and various biotic factors, can significantly impair cucumber growth by disrupting physiological processes [89, 90]. Excessive water impairs oxygen supply to the root system, leading to varying degrees of damage. One of the primary phenotypic responses of cucumber to waterlogging stress is the formation of adventitious roots (AR) (Figure 8). Adventitious roots (ARs) and lateral roots (LRs) originate from stem and root tissues, respectively, playing an essential function in the development processes of plants [91].

The formation of adventitious roots (AR) is largely governed by hormonal regulation [92]. Auxin is crucial for regulating root growth in cucumber plants, from controlling directional growth to promoting the development of lateral and adventitious roots [93]. Its influence is essential at various stages of root formation [94]. Several researches have emphasized the function of auxin in regulation of adventitious roots (ARs) and lateral roots (LRs) development in cucumber. For example, the concentration of endogenous auxin in the hypocotyl was elevated 72 hours following exposure to waterlogging stress. Additionally, the application of 10 mg/L NAA further promoted the formation of adventitious roots (AR) [92]. The present study investigates the impact of waterlogging stress and its interaction with exogenous hormones on the phenotype of *Cucumis sativus* hypocotyls. After 48 hours of waterlogging (WL) stress, adventitious root (AR) formation was significantly impaired compared to controls, consistent with previous reports highlighting the inhibitory effects of waterlogging on root development [93, 95]. However, when WL stress was combined with NAA or ETH, a notable increase in AR formation was observed, suggesting that these hormones may mitigate the adverse effects of waterlogging on root growth [28, 96]. In contrast, no ARs were observed in plants treated with waterlogging plus MeJA, which aligns with findings that MeJA can inhibit root formation under stress conditions [93]. Interestingly, after four days, WL combined with NAA promoted AR formation to a greater extent than WL with ETH, indicating that NAA may be more effective in stimulating root development under waterlogging stress [96]. These results support the hypothesis that exogenous hormones, particularly NAA and ETH, can alleviate waterlogging-induced root inhibition, while MeJA may exacerbate stress responses. Additional research is necessary to understand the molecular mechanisms governing these hormonal interactions in cucumber.

Subsequent investigation demonstrated that the application of auxin stimulated the expression of genes responsible for ethylene biosynthesis (*CsACS1*, *CsACS2*, *CsACO5*) as well as genes associated with reactive oxygen species (ROS) signaling pathways, such as CsRBOHB and CsRBOHF3, in response to waterlogging stress [92]. In contrast, jasmonate, also known MeJA treatment significantly impaired the formation of adventitious roots (AR) [93]. A previous study also showed that JA negatively affects AR development. In the waterlogging-sensitive Pepino line, AR growth was inhibited due to elevated endogenous JA levels, whereas the resistant Zaoer N line exhibited lower JA accumulation [93, 97]. In this study, MeJA application also inhibited AR growth under waterlogging stress. However, the underlying mechanism by which JA restricts adventitious root development under such conditions requires further investigation.

Under waterlogging stress, *CsC3H9* and *CsC3H13* exhibited a significant increase in expression, suggesting their involvement in stress response mechanisms, which aligns with prior studies indicating the function of C3H genes in abiotic stress tolerance [12]. In contrast, other *CsC3H* genes, including *CsC3H2*, *CsC3H4*, *CsC3H7*, *CsC3H11*, *CsC3H12*, and *CsC3H15*, were down-regulated, reflecting a selective activation of specific family members under hypoxic and oxidative stress conditions caused by waterlogging. Hormonal treatments modulated CsC3H gene expression, with NAA significantly upregulating *CsC3H2*, *CsC3H3*, *CsC3H12*, *CsC3H13*, and *CsC3H14*, which are involved in adventitious root (AR) formation under stress [98]. Conversely, *CsC3H4*, *CsC3H7*, *CsC3H11*, and *CsC3H15* showed reduced expression, suggesting a potential negative regulatory role in AR formation. Ethylene (ETH) treatment induced a strong response in *CsC3H6*, *CsC3H12*, and *CsC3H15*, confirming ethylene’s known role in regulating stress tolerance [99]. In contrast, MeJA treatment down-regulated most *CsC3H* genes, except for *CsC3H5*, *CsC3H7*, and *CsC3H11*, indicating their specific response to jasmonic acid signaling during waterlogging stress [9]. *CsC3H6* consistently upregulated across all treatments, suggesting its role as a pleiotropic regulator that integrates multiple stress and hormonal signals [9]. These findings highlight *CsC3H* genes, particularly *CsC3H6*, *CsC3H9*, *CsC3H12*, and *CsC3H13*, as key players in cucumber’s response to waterlogging stress and hormone signaling, with potential applications in improving stress tolerance in breeding programs.

In addition, we investigated the tissue-specific expression patterns of *CsC3H* genes, given the well-documented function of the C3H gene family in modulating plant development and growth, as emphasized in a range of scholarly publications. There were slight differences in the expression profiles of *CsC3H* genes across various cucumber tissues when comparing microarray and qRT-PCR analyses. These discrepancies could be due to factors such as variations in sampling time, genetic differences, or the methodologies employed in the experiments. The expression levels of C3H genes in cucumber exhibited notable variation across different tissues and developmental stages [4, 9, 15]. Specifically, the expression of *CsC3H9* and *CsC3H14* was upregulated in female flowers, while *CsC3H15* was expressed in male flowers.

Notably, *CsC3H9* and *CsC3H15* showed flower-specific expression, suggesting their potential involvement in regulating flowering in cucumber. *CsC3H5, CsC3H22*, and *CsC3H25* were detected in leaf, root, and stem tissues but not in tendrils, indicating a possible role in leaf development. Furthermore, two genes were detected in the root tissue, while one gene was found in the ovary. The expression profiles of C3H genes in cucumber exhibit distinct variations compared to those in other species, including pepper, Arabidopsis, and rice, where the majority of C3H genes are predominantly expressed in the roots, flowers, foliage, and seeds [9, 66]. These tissue-specific expression patterns of CsC3H genes provide valuable insights into their potential roles in regulating various developmental processes in cucumber, particularly in the formation of flowers, roots, and leaves. Further functional validation of these genes will be crucial for understanding their precise contributions to cucumber growth and development. Proteins required to be localized to the appropriate cellular compartment in order to function effectively. A protein’s subcellular location can provide valuable insights into its potential role and function [100, 101]. CCCH-type zinc-finger proteins display a variety of localization patterns within the cell. A subset of these proteins, such as *GhZFP1*, is primarily localized to the nucleus [102], *AtTZF11* [103], *OsDOS* [104], and *AtZFP1* [105]. Certain proteins are confined to specific cellular compartments, including the cytoplasm or plasma membrane, as exemplified by Oxidation-related Zinc Finger 1 (*AtOZF1*) [106] and *AtOZF2* [107]. Whereas some CCCH zinc-finger proteins are found in the cytoplasm, including *ZFP36L3* [108] and *AtTZF2/3* [16], while others, such as *AtTZF7* and *OsLIC* [74, 109], can shift between the nucleus and the cytoplasm. Our research revealed that the GFP-tagged *CsC3H9* was exclusively localized to the nucleus (Figure 12), suggesting its potential involvement in modulating transcription factor function and influencing the outcomes of target proteins. We hypothesize that cucumber C3H proteins might collectively participate in the plant’s response to waterlogging stress through cooperative interactions among different members of the family. Further studies are required to delineate the precise network of C3H protein interactions involved in regulating waterlogging stress, in order to clarify the contribution of C3H transcription factors within this context.

In summary, we present a schematic model outlining the key *CsC3H* genes associated in response of *C. sativus* to waterlogging stress and various exogenous hormone treatments (Fig. 13). This study provides an in-depth investigation of the functional roles and expression profiles of the *CsC3H* gene family in cucumber. Our results offer crucial understanding into the molecular mechanisms that regulate the cucumber’s response to waterlogging stress, forming a basis for future research in this area.

**Fig. 13.**
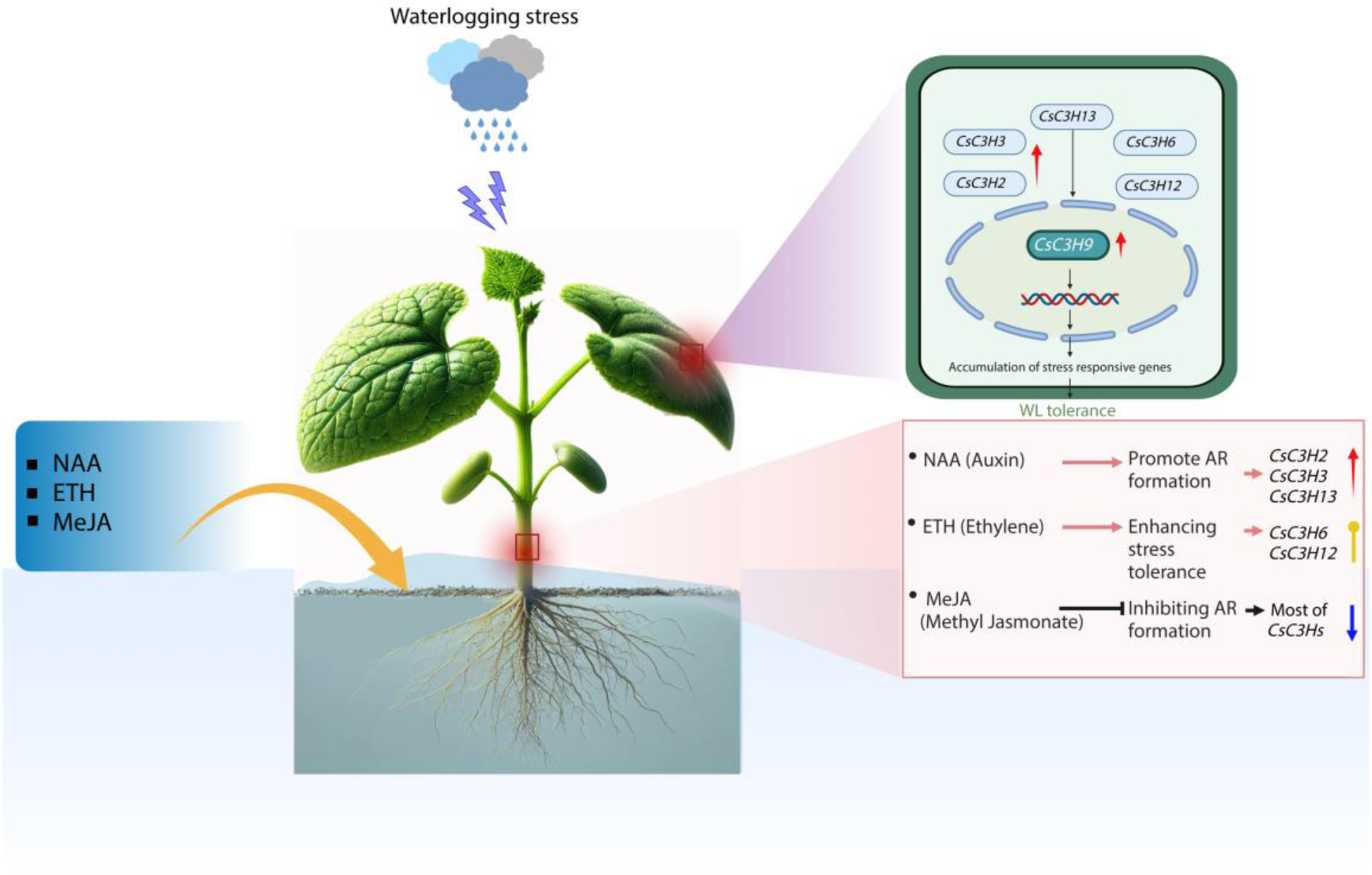
A schematic model illustrating the subcellular distribution of C3H proteins and the transcriptional patterns of *CsC3H* transporters in the leaves and roots of *C. sativus* in waterlogging and exogenous hormone stress is presented. The model highlights the following: the red arrows represent significant overexpression of the genes, the blue arrows indicate marked downregulation, and the yellow arrows denote the positive activation of gene expression changes.

## 5. Conclusion

The first extensive genome-wide identification of C3H genes in cucumber, identifying and characterizing 38 *CsC3H* genes within the cucumber genome. Through detailed analyses of phylogenetic relationships, chromosomal distribution, gene duplication events, selective evolutionary pressures, conserved domains, gene structure, and cis-regulatory elements, the functional diversity of *CsC3H* genes was revealed. Additionally, quantitative PCR analysis of 12 *CsC3H* genes suggests their significant role in cucumber’s response to hormones and waterlogging stress. The transcriptional expression profile of *CsC3H* genes across various tissues, and under treatments with NAA, ETH, and MeJA, were examined. Subcellular localization studies confirmed that GFP-tagged *CsC3H9* is specifically localized in the nucleus. Our results contribute to a deeper understanding of the *CsC3H* gene family in cucumber, providing critical insights into the role of *CsC3H* proteins in mediating responses to hormonal signals and waterlogging stress. These findings have significant implications for the development of future cucumber breeding strategies.

